# Out of the blue: Family-wide loss of anthocyanin biosynthesis in Cucurbitaceae

**DOI:** 10.1101/2025.10.06.680802

**Authors:** Nancy Choudhary, Marie Hagedorn, Boas Pucker

## Abstract

- Plant pigmentation secrets are among the oldest interests of plant scientists, with pigments such as chlorophyll, carotenoids, flavonoids, and betalains contributing to the diversity of hues in higher plants. Anthocyanins, a class of flavonoids responsible for vibrant shades of pink, red, and blue pigmentation, are almost ubiquitous in angiosperms but are replaced by betalains in some families in the order Caryophyllales.
- We investigated anthocyanin pigmentation in Cucurbitaceae, one of the largest fruit and vegetable families, characterised by white and yellow flowers and red, orange, and green fruits predominantly pigmented by carotenoids. Using a comprehensive collection of 258 datasets representing 183 unique species across all 15 tribes of Cucurbitaceae, along with a phylogenomics approach, we observed a systematic absence of genes involved in anthocyanin and proanthocyanidin biosynthesis. Absence of the structural genes *DFR*, *ANS*, *arGST*, *LAR*, and *ANR*, along with the anthocyanin-related regulatory MYB genes, was consistently confirmed by synteny and phylogenetic analysis.
- These results suggest that anthocyanin loss in angiosperms is more common than previously assumed. In light of this new discovery, we propose a stepwise loss of anthocyanin pigmentation in Cucurbitaceae, likely accompanied by a partial functional replacement with carotenoid pigmentation.

## Introduction

The colours of plants are one of the most striking manifestations of plant evolution, arising from and reinforcing interactions with the environment through pollinator attraction, seed dispersal, defence and visual signalling (Lev-Yadun *et al*., 2004; Chittka & Raine, 2006; Lev-Yadun, 2016; Garcia *et al*., 2019). Dawned with complex modifications, anthocyanins, derived from the flavonoid pathway, are one of the key contributors to this chromatic diversity in flowering plants. Widely distributed in seed plants, they are primarily responsible for the diversity of hues seen in most plant tissues, specifically the vibrant red, pink, and blue hues of flowers and fruits (Winkel-Shirley, 2001). Beyond aesthetics, they serve multiple ecological and physiological functions (Grünig *et al*., 2025), including photoprotection against ultraviolet (UV) radiation (Gould, 2004), attraction of pollinators and seed dispersers (Thompson *et al*., 1972), and defence responses against herbivory and pathogen attack (Steyn *et al*., 2002). Moreover, anthocyanins have been implicated in plant adaptation to environmental stress, including drought and cold tolerance (Chalker-Scott, 1999; Pietrini *et al*., 2002).

The core flavonoid biosynthesis pathway is well understood and begins with the synthesis of naringenin chalcone from one molecule of 4-coumaroyl-CoA and three molecules of malonyl-CoA, catalysed by chalcone synthase (CHS) (**Fig. 1a**). Naringenin chalcone is rapidly and stereospecifically isomerised to the colourless naringenin by chalcone isomerase (CHI). From this central intermediate, multiple enzymatic steps branch into distinct flavonoid classes. Naringenin can be directly converted into flavones by flavone synthase (FNS). Alternatively, flavanone 3-hydroxylase (F3H) catalyses the conversion of naringenin into dihydrokaempferol, which undergoes B-ring hydroxylation by flavonoid 3′-hydroxylase (F3′H) and flavonoid 3′,5′-hydroxylase (F3′5′H), producing structurally diverse dihydroflavonols. These dihydroflavonols are then converted by flavonol synthase (FLS) into flavonols or channelled towards anthocyanin production via dihydroflavonol reductase (DFR), anthocyanidin synthase (ANS), and anthocyanin-related glutathione S-transferase (arGST). Unstable anthocyanidins are converted to stable anthocyanins by UDP-glucose flavonoid/anthocyanidin 3-O-glycosyltransferase (UGT). Glycosylation, acylation, and methylation further modify the properties of the anthocyanidin-3-glucoside derivatives and generate a variety of anthocyanins. Synthesised on the cytosolic face of the endoplasmic reticulum, anthocyanins must be sequestered into the vacuole for stabilisation and accumulation. Two major mechanisms have been proposed: (i) GST-mediated ligand transport, in which glutathione S-transferases act as carrier proteins to facilitate anthocyanin transport through ABCC or MATE-type transporters (Mueller *et al*., 2000; Sun *et al*., 2012), and (ii) vesicle-mediated trafficking, where anthocyanins are packaged into vesicles and delivered to the vacuole (Poustka *et al*., 2007) (**Fig. 1b**). In addition to anthocyanins, the flavonoid pathway produces proanthocyanidins (condensed tannins), which play crucial roles in plant defence, particularly through anti-herbivory and anti-pathogen activities, and contribute to seed coat pigmentation and dormancy (Xie *et al*., 2003; Dixon *et al*., 2005). The enzymatic steps diverging from the general flavonoid pathway, mediated by leucoanthocyanidin reductase (LAR) and anthocyanidin reductase (ANR), direct metabolic flux towards proanthocyanidin biosynthesis (**Fig. 1a**) (Dixon *et al*., 2005; Lepiniec *et al*., 2006).

**Fig. 1:**
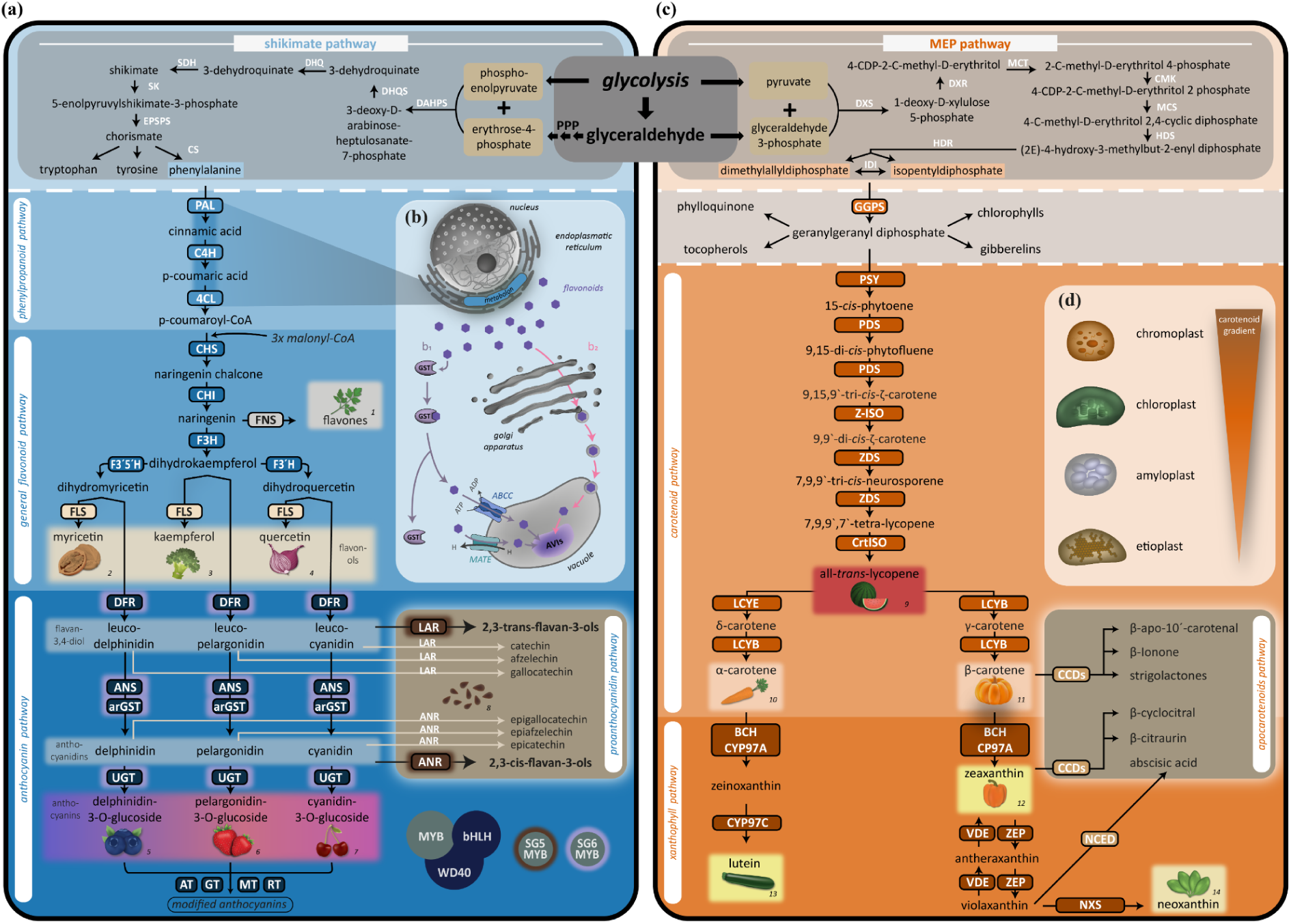
Key enzymes of the two major pigmentation pathways in plants: (a) flavonoid biosynthesis pathway and (c) carotenoid biosynthesis pathway. (a) Flavonoid biosynthesis begins with phenylalanine, which is synthesised via the shikimate pathway. Branches of this pathway, including the anthocyanin and proanthocyanidin pathways, contribute strongly to visible pigmentation in higher plants. DAHPS, 3-deoxy-D-arabino-heptulosonate-7-phosphate synthase; DHQS, 3-dehydroquinate synthase; DHQ, 3-dehydroquinate dehydratase; SDH, shikimate 5-dehydrogenase; SK, shikimate kinase; EPSPS, 5-enolpyruvylshikimate 3-phosphate synthase; CS, chorismate synthase; PAL, phenylalanine ammonia-lyase; C4H, cinnamate 4-hydroxylase; 4CL, 4-coumarate-CoA ligase; CHS, chalcone synthase; CHI, chalcone isomerase; FNS, flavone synthase; F3H, flavanone 3-hydroxylase; F3′H, flavonoid 3′-hydroxylase; F3′5′H, flavonoid 3′,5′-hydroxylase; FLS, flavonol synthase; DFR, dihydroflavonol 4-reductase; ANS, anthocyanidin synthase; arGST, anthocyanin-related glutathione S-transferase; UGT, UDP-glycosyltransferase; AT, acyltransferase; GT, glycosyltransferase; MT, methyltransferase; RT, rhamnosyltransferase; LAR, leucoanthocyanidin reductase; ANR, anthocyanidin reductase. (b) Flavonoids are synthesised at the endoplasmic reticulum and subsequently transported into the vacuole for further modifications and storage. Current evidence supports two major transport mechanisms: ligand-based transport (b1) and vesicle-mediated transport (b2). GST, glutathione S-transferase; ABCC, ATP-binding cassette subfamily C; MATE, multidrug and toxic compound extrusion; AVI, anthocyanin vacuolar inclusions. (c) Carotenoid biosynthesis is initiated by the condensation of isopentenyl diphosphate and dimethylallyl diphosphate, derived from the MEP (2-C-methyl-D-erythritol 4-phosphate) pathway. The first committed step is catalysed by PSY, which converts geranylgeranyl diphosphate to 15-cis-phytoene. Following sequential desaturation and isomerisation steps, lycopene is formed, which serves as the branching point into the α-carotene and β-carotene lineages. Carotenoids are further modified to generate diverse xanthophylls, and cleavage by CCDs or NCEDs produces apocarotenoids. DXS, 1-deoxy-D-xylulose 5-phosphate synthase; DXR, DXP reductoisomerase; MCT, MEP cytidyltransferase; CMK, 4-(cytidine 5′-diphospho)-2-C-methyl-D-erythritol kinase; MCS, 2-C-methyl-D-erythritol 2,4-cyclodiphosphate synthase; HDS, 1-hydroxy-2-methyl-2-butenyl 4-diphosphate synthase; HDR, HMBPP reductase; IDI, isopentenyl diphosphate isomerase; GGPS, geranylgeranyl diphosphate synthase; PSY, phytoene synthase; PDS, phytoene desaturase; Z-ISO, ζ-carotene isomerase; ZDS, ζ-carotene desaturase; CrtISO, carotenoid isomerase; LCYE, lycopene ε-cyclase; LCYB, lycopene β-cyclase; CYP97A/C, cytochrome P450-type monooxygenases 97A and 97C; BCH, β-carotene hydroxylase; VDE, violaxanthin de-epoxidase; ZEP, zeaxanthin epoxidase; NXS, neoxanthin synthase; CCDs, carotenoid cleavage dioxygenases; NCED, 9-cis-epoxycarotenoid dioxygenase. Carotenoid biosynthesis and storage occur in plastids. (d) The accumulation of carotenoids in different plastid types is shown. The fruit and vegetable icons illustrate examples of known plants enriched in specific pigments (1 parsley, 2 walnut, 3 broccoli, 4 red onion, 5 blueberry, 6 strawberry, 7 cherry, 8 apple seeds, 9 watermelon, 10 carrot, 11 pumpkin, 12 orange bell pepper, 13 zucchini, 14 spinach).

The regulation of anthocyanidin and proanthocyanidin biosynthesis is tightly controlled by the MBW transcriptional complex, which is composed of MYB (myeloblastosis), bHLH (basic helix-loop-helix), and WD40 proteins (Koes *et al*., 2005; Jaakola, 2013). Well-characterised MBW complex examples include the TRANSPARENT TESTA2, TRANSPARENT TESTA8, and TRANSPARENT TESTA GLABRA1 (*At*TT2-*At*TT8-*At*TTG1) complex in *Arabidopsis*, which regulates proanthocyanidin biosynthesis (Gonzalez *et al*., 2009) and the ANTHOCYANIN2 (*Ph*AN2)-*Ph*AN1-*Ph*AN11 complex in *Petunia,* regulating anthocyanin production (Spelt *et al*., 2000). Certain R2-R3 MYB proteins involved in flavonoid biosynthesis generally act as the primary activators of branch-specific genes and determine specificity, while bHLH and WD40 proteins provide stabilisation and combinatorial control (Gonzalez *et al*., 2008; Hichri *et al*., 2011). The flavonoid-related WD40 and bHLH proteins are pleiotropic and involved in multiple processes beyond anthocyanin synthesis (Walker *et al*., 1999; Spelt *et al*., 2002; Bernhardt *et al*., 2003; Airoldi *et al*., 2019).

Pigmentation in plants is not limited to flavonoids. Chlorophylls, the most abundant pigments, confer green colouration and are essential for photosynthesis. Carotenoids, typically yellow, orange, or red, are plastid-derived isoprenoids produced via the methylerythritol phosphate (MEP) pathway (**Fig. 1c**). Phytoene synthase (PSY) catalyses the first committed step, condensing geranylgeranyl diphosphate (GGPP) molecules into phytoene. Sequential desaturation and isomerisation steps mediated by phytoene desaturase (PDS), ζ-carotene desaturase (ZDS), and carotenoid isomerases yield lycopene, the central branching point. Cyclisation reactions then give rise to the α- and β-carotene branches, which are further modified by hydroxylases and epoxidases to synthesise lutein, zeaxanthin, violaxanthin, and neoxanthin (Cazzonelli & Pogson, 2010). Unlike anthocyanins, carotenoids are primary metabolites and are essential for plant life. They are involved in photoprotection and light harvesting (Triantaphylidès & Havaux, 2009), act as precursors of signalling molecules like abscisic acid and strigolactones, and accumulate in chromoplasts (**Fig. 1d**) to attract pollinators and seed dispersers (Sun *et al*., 2022).

In plants, another class of pigments, betalains, occur exclusively in the order Caryophyllales (Mabry & Mabry, 1964). Derived from tyrosine, they produce red, yellow, and violet colours and have completely replaced anthocyanins in multiple lineages within this order (Stafford, 1994; Brockington *et al*., 2011; Sheehan *et al*., 2020). This mutual exclusivity, potentially resulting from the loss of arGST and MBW complex members, demonstrates dynamic shifts in pigmentation pathways during evolution (Timoneda *et al*., 2019; Pucker *et al*., 2024). By contrast, anthocyanins and carotenoids often coexist within the same species, organs, or even the same tissues, increasing colour variety (Van Eijk *et al*., 1987; Ooe *et al*., 2016; Clemente *et al*., 2023).

Although loss of anthocyanin pigmentation has been reported, such cases are typically restricted to individual species or even cultivar-level variants, often caused by single-gene mutations or regulatory changes (Smith & Goldberg, 2015; Marin-Recinos & Pucker, 2024). Large-scale, lineage-wide loss is exceedingly rare. One notable exception is the genus *Brachypodium*, where all examined species lack key anthocyanin biosynthetic genes, including MYB regulators, *ANS*, and *arGST* (Khatun *et al*., 2025). In the grass family (*Poaceae*), canonical anthocyanin MYBs and TT19-type arGSTs have been lost and functionally replaced by non-canonical anthocyanin MYBs and BZ2-type arGSTs, preserving anthocyanin production via an alternative molecular architecture (Khatun *et al*., 2025).

The *Cucurbitaceae* family (also called cucurbits), which consists mainly of tropical and subtropical distribution, herbaceous annual or perennial climbers and woody lianas, were among the first plants cultivated by humans (Bisognin, 2002; Schaefer *et al*., 2008). Harbouring economically important fruit and vegetable crops like pumpkins, gourds, melons, cucumbers, and squashes, the family contains ∼115 genera and ∼1000 species. Their fruit and flower colour ranges from green, yellow, and orange to various shades of red (Schaefer & Renner, 2011a). Historically, carotenoids have been associated with yellow and orange pigmentation in cucurbits (Azevedo-Meleiro & Rodriguez-Amaya, 2007), while chlorophylls explain green tissues. To the best of our knowledge, blue or purple pigments have not been documented in the Cucurbitaceae; while comprehensive analyses of pigment diversity are lacking, current data offer no reason to expect their presence.

In this study, we performed a comprehensive phylogenomic analysis to elucidate the genetic basis of pigmentation in the Cucurbitaceae family. Leveraging publicly available genomic and transcriptomic resources, we systematically surveyed over 180 species, representing all 15 tribes of Cucurbitaceae, for the presence of key structural and regulatory genes involved in flavonoid biosynthesis. Our results reveal a consistent absence of essential anthocyanin biosynthetic genes, including the structural and regulatory genes. Additionally, we identified parallel losses in the proanthocyanidin branch genes. This widespread loss of core flavonoid biosynthesis components strongly suggests an evolutionary abandonment of anthocyanin and proanthocyanidin pigmentation pathways within Cucurbitaceae and highlights carotenoids as the primary alternative. Our findings provide key insights into the evolutionary remodelling of pigmentation systems and lay a foundation for understanding the molecular determinants of colour diversity in cucurbits.

## Material and Methods

### Data collection, annotation, and preparation of final datasets

The species analysed in this study include members of Cucurbitales, primarily from Cucurbitaceae, with outgroups from Datiscaceae, Begoniaceae, and Coriariaceae, as well as representatives from the orders Rosales and Fagales. Sequence datasets were retrieved from the National Center for Biotechnology Information (NCBI; https://www.ncbi.nlm.nih.gov/, last accessed July 2025), Cucurbit Genomics Database (CucGenDBv2; http://cucurbitgenomics.org/v2/), Watermelon Genomics and Mutation Database (WaGMDB; http://www.watermelondb.cn), Melonomics v4 (http://melonomics.net), GigaDB, Figshare, Genome Warehouse (GWH; https://ngdc.cncb.ac.cn/gwh/), and the China National GeneBank Sequence Archive (CNSA; https://db.cngb.org/cnsa). In total, 282 genomic and transcriptomic datasets were obtained, comprising 258 *Cucurbitaceae* datasets and additional outgroups (for data sources, see Dataset_sources.csv (Choudhary *et al*., 2025)). For the transcriptomic datasets, only coding sequences were available and downloaded directly.

Eighty-two genome assemblies from NCBI lacked corresponding annotations and were annotated using GeMoMa v1.9 (Keilwagen *et al*., 2016). This was performed with reference to high-quality (96.7-99.8 % BUSCO conservation) annotations of ten species: *Alnus glutinosa* (Christenhusz *et al*., 2024), *Juglans regia* (Marrano *et al*., 2020), *Quercus robur* (GCA_932294415.1), *Sicyos edulis* (Fu *et al*., 2021), *Benincasa hispida* (Xie *et al*., 2019), *Cucumis sativus* (Li *et al*., 2019), *Cucumis melo* (GCA_025177605.1), *Cucurbita maxima* (Sun *et al*., 2017), *Cucurbita pepo* (Montero-Pau *et al*., 2018), and *Momordica charantia* (Urasaki *et al*., 2017). For each annotated species, the ten GeMoMa outputs were merged and filtered using the parameters f=“start==’M’ and stop==’*’ and (isNaN(score) or score/aa>=’0.75’)” and atf=“sumWeight>5”. To assess the completeness of available annotations, an additional seven species were re-annotated using the same parameters described above using GeMoMa v1.9 (For annotation details, see Gene_annotation_stats.csv (Choudhary *et al*., 2025)). The gene annotation datasets are publicly available via bonndata (Choudhary *et al*., 2025). For all genomic datasets, genome sequence (.fasta) and annotation (.gff3) files were retrieved, and coding sequences (CDS) were extracted using agat_sp_extract_sequences.pl from the AGAT toolkit (Dainat, 2024). All CDS were retained for flavonoid and carotenoid biosynthesis gene identification and expression analysis. To streamline downstream analyses and reduce redundancy, only the longest transcript per gene was selected for use in gene trees, species tree, synteny analysis, and whole-genome duplication (WGD) detection. For this, agat_sp_keep_longest_isoform.pl was used. CDS were translated to protein sequences using a custom Python script ‘transeq.py’ v0.3 available at https://github.com/bpucker/PBBtools.

The completeness of all protein datasets was assessed using BUSCO v5.8.1 (Tegenfeldt *et al*., 2025) in protein mode with the eudicotyledons_odb12 reference dataset. Since some Cucurbitaceae datasets were found to be contaminated (see **Fig. S15**), to systematically address such cases and ensure consistent quality of datasets, all Cucurbitaceae CDS were taxonomically classified using Kraken v2.1.3 (Wood *et al*., 2019) with the ‘core_nt’ database (December 2024 release, containing GenBank, RefSeq, TPA and PDB entries). Bracken v2.9 (Lu *et al*., 2017) was applied to the Kraken outputs to estimate species-level abundance. Any dataset with >10% of classified sequences assigned to a non-Cucurbitaceae taxon was flagged as “contaminated” (for detailed classification report, see Kraken_classification.csv (Choudhary *et al*., 2025)). These contaminated datasets were excluded from species and gene phylogenies and no conclusions were based on these datasets.

### Species phylogeny and flower and fruit colour mapping

All non-contaminated datasets were used to construct the species phylogeny. Single-copy orthologs were identified using BUSCO_phylogenomics.py (https://github.com/jamiemcg/BUSCO_phylogenomics/) across 2805 unique BUSCO-defined single-copy marker genes (*eudicotyledons_odb12*). Peptide sequences of each ortholog group were aligned with MUSCLE v5.3 (Edgar, 2022), and poorly aligned regions were trimmed with trimAl v1.5.rev0 (Capella-Gutiérrez *et al*., 2009) using the -automated1 option. For the first tree, a multi-species coalescent approach was used, where individual gene trees were inferred for all 2805 alignments using FastTree v2.1.11 (Price *et al*., 2010) under the default JTT+CAT model, followed by species tree estimation using wASTRAL v1.23.3.7 (-R -u 2) (Zhang & Mirarab, 2022). Based on this initial tree, five datasets were removed due to their suspicious phylogenetic placement (see results section for details). For the final species phylogeny, both multi-species coalescent and concatenation (supermatrix) approaches were used to infer the Cucurbitaceae phylogeny. For the coalescent-based phylogeny, 20 of the 2805 alignments were randomly selected and individually tested with ModelFinder (Kalyaanamoorthy *et al*., 2017), which most frequently identified the Q.plant+I+G4 model. This model was therefore applied to all gene trees for consistency. Gene trees were inferred using IQ-TREE v2.3.6 (Minh *et al*., 2020) with the parameters --bb 1000 --alrt 1000 -m Q.plant+I+G4. The species tree was estimated from the gene trees using wASTRAL v1.23.3.7 (Zhang & Mirarab, 2022) using -R and -u 2 parameters. For the concatenation-based species phylogeny, only BUSCO orthogroups present in at least 80% of species were used for the supermatrix construction, leading to the use of 516 BUSCO genes. All trimmed alignments for these orthogroups were concatenated into a supermatrix within IQ-TREE v2.3.6 (Minh *et al*., 2020), followed by model selection with ModelFinder (Kalyaanamoorthy *et al*., 2017) and tree inference. BUSCO genes used for coalescence- and concatenation-based species phylogeny can be found in BUSCO_genes_for_species_phylogeny.csv (Choudhary *et al*., 2025). Fruit and flower colour information was obtained from published literature, floras, and species descriptions (Fruit_flower_colour_sources.csv (Choudhary *et al*., 2025)).

### Identification of flavonoid and carotenoid biosynthesis genes

KIPEs3 v0.38 (Rempel *et al*., 2023) was used to identify the structural genes in the carotenoid and flavonoid biosynthesis pathways, utilising the polypeptide sequences of all the analysed datasets. The provided baits and residues datasets (FlavonoidBioSynBaits_v3.4 and CarotenoidBioSynBaits_v2.0; available at https://github.com/bpucker/KIPEs) were used to detect and functionally assess candidate sequences. KIPEs3 identifies homologs through sequence similarity to curated bait sequences and evaluates their functional integrity by comparing the conservation of experimentally verified, functionally important amino acid residues for each enzyme. The output includes the percentage of conserved functional residues, which serves as an indicator of potential enzymatic activity and was also used in subsequent analysis and visualisations.

For transcriptional regulators, members of the MBW complex were identified as follows: MYB candidates using MYB annotator v0.2 (Pucker, 2022), bHLH candidates using bHLH annotator v0.2 (Thoben & Pucker, 2023), and WD40 candidates by constructing phylogenetic trees including previously characterised anthocyanin-regulating WD40 sequences from the literature. Phylogenetic trees were also generated as an additional line of evidence for confirming the presence/absence of all identified genes (**Fig. S4-12**). Structural gene candidates predicted by KIPEs3 that retained at least 75% of residues known to be critical for function were retained to ensure that even potential pseudogenes are captured when inferring gene loss. Previously characterised functional sequences from the literature were also included (see Bait_sequences_gene_trees.csv for sequence IDs (Choudhary *et al*., 2025)). Multiple sequence alignments were generated separately for the following groups: CHS, CHI, CYP450 (cytochrome P450, including FNS II, F3′H, and F3′5′H), 2ODDs (2-oxoglutarate-dependent dioxygenases, including FNS I, F3H, FLS, and ANS), SDRs (including DFR, LAR, and ANR), arGST, UGTs (including F3GT and A3GT), MYBs (SG5, SG6, and SG7 MYBs), anthocyanin-associated bHLHs, and anthocyanin-associated WD40 proteins.

For the regulatory gene candidates, the script collect_best_blast_hits.py (available at https://github.com/bpucker/ApiaceaeFNS1) (Pucker & Iorizzo, 2023) was used to collect similar sequences to known MBW genes from closely-related *Malus domestica* and the model plant *Arabidopsis thaliana*. These sequences, together with reference MBW sequences from the literature, were used for phylogenetic analysis. Coding sequences used for all gene trees are publicly available (Choudhary *et al*., 2025). Alignments were generated using MAFFT v7.490 (G-INS-i algorithm) (Katoh *et al*., 2019) and translated back to codon alignments with pxaa2cdn (Brown *et al*., 2017). Alignments were filtered with pxclsq from the PHYX toolkit (-p 0.1) (Brown *et al*., 2017). Maximum-likelihood phylogenies were inferred using IQ-TREE v2.3.6 (Minh *et al*., 2020) with the parameters: --alrt 1000 -B 1000 -m MFP, where the best-fit evolutionary model was determined by ModelFinder (Kalyaanamoorthy *et al*., 2017).

### Synteny analysis and WGD dating

Genome-wide syntenic and collinear blocks were identified across 19 representative genome sequences included in this study; *Fragaria vesca*, *Ulmus minor*, *Quercus robur*, *Coriaria nepalensis*, *Begonia loranthoides*, *Begonia peltatifolia*, *Bayabusua clarkie*, *Gynostemma pentaphyllum*2, *Thladiantha cordifolia*1, *Momordica charantia*2, *Herpetospermum pedunculosum*1, *Luffa aegyptiaca*1, *Sicyos edulis*1, *Trichosanthes truncata*1, *Cucurbita moschata*1, *Cucurbita pepo*1, *Citrullus lanatus*2, *Cucumis melo*1, and *Cucumis sativus*1. These genome sequences were selected based on completeness (as measured by the BUSCO score), continuity, and taxonomic diversity. Synteny analysis was performed using JCVI/MCscan v1.4.14 following the package workflow:

https://github.com/tanghaibao/jcvi/wiki/Mcscan-(python-version). Coding sequence (CDS) and GFF annotation files were used as input. Pairwise orthology and synteny searches were conducted using *F. vesca* as the reference, with default parameters and a c-score cutoff of 0.99. For each species, synteny depth was plotted against *F. vesca* to determine the appropriate number of syntenic block iterations, which is critical for accurately representing synteny in species with whole-genome duplications (WGDs). Based on this approach, the inferred synteny depth ratios against one region of *F. vesca* were as follows: *U. minor*:1, *Q. robur*:1; *C. nepalensis*:4, *B. loranthoides*:4, *B. peltatifolia*:4, *B. clarkei*:2, *G. pentaphyllum*:2, *T. cordifolia*:2, *M. charantia*:2, *H. pedunculosum*:2, *L. aegyptiaca*:2, *S. edulis*:4, *T. truncata*:2, *C. moschata*:4, *C. pepo*:4, *C. lanatus*:2, *C. melo*:2, *C. sativus*:2.

To investigate *DFR*, *ANS*, and *arGST* synteny, candidate sequences were identified in *F. vesca* and used to extract conserved syntenic regions across all selected species. For *arGST*, the flanking genomic regions were less conserved than for *DFR* and *ANS*, possibly due to genome rearrangements associated with speciation and WGDs. To ensure robust orthology assignment, we used a phylogenomics approach, consisting of (i) extraction of ±35 flanking genes around the *F. vesca arGST* locus, (ii) identification of candidate homologs using collect_best_blast_hits.py (available at https://github.com/bpucker/ApiaceaeFNS1) (Pucker & Iorizzo, 2023), (iii) multiple sequence alignment using MAFFT v7.490 (--auto strategy) (Katoh *et al*., 2019), (iv) alignment cleaning with pxclsq (Brown *et al*., 2017), and (v) phylogenetic inference using FastTree v2.1.11 (Price *et al*., 2010), under default parameters. Only the syntelogs (orthologs in a macrosyntenic context) were accepted as syntenic orthologs and used for downstream visualisation.

Since synteny depth suggested four syntenic regions of *Coriaria* against one region of *F. vesca*, this suggested a previously unreported WGD event in *C. nepalensis*. To test this, WGD analysis was performed using wgd v2.0.38 (Chen & Zwaenepoel, 2023; Chen *et al*., 2024) based on coding sequences of high-quality genome sequences (*Q. robur*, *C. nepalensis*, *B. loranthoides*, *G. pentaphyllum*2, and *C. sativus*1). Default workflows were followed to infer orthogroups, compute synonymous substitutions per site of duplicated genes (Ks distributions), detect intra-and interspecies synteny, and estimate absolute WGD ages via Bayesian MCMC dating. The species phylogeny used for dating was reconstructed under a multi-species coalescent framework as described above. The following lower and upper-bound divergence times were used as secondary calibration points from timetree.org (Kumar *et al*., 2022): *Q. robur* vs. Cucurbitales (93.0-109.4 mya); *C. nepalensis* vs. *B. loranthoides* (55.8-76.0 mya); *B. loranthoides* vs. Cucurbitaceae (56.6-91.0 mya); and *G. pentaphyllum* vs. *C. sativus* (26.5-68.7 mya).

### Expression analysis

To compare the expression of remaining flavonoid biosynthesis genes in Cucurbitaceae with those of closely related outgroups, we collected and analysed publicly available RNA-seq datasets from NCBI Sequence Read Archive (SRA, https://www.ncbi.nlm.nih.gov/sra/) following a previously established approach (Pucker *et al*., 2024). Species were selected based on three criteria: (i) broad taxonomic representation, (ii) high completeness of gene annotation (protein BUSCO), and (iii) availability of >10 RNA-seq datasets to ensure robust comparisons.

All paired-end RNA-seq FASTQ files available at NCBI SRA were collected for all selected species using the NCBI SRA toolkit v3.1.0 (available at https://github.com/ncbi/sra-tools). Transcript abundance was quantified using kallisto v0.50.1 (Bray *et al*., 2016) based on the coding sequences. The individual count tables were then merged and filtered with customised Python scripts (Pucker & Iorizzo, 2023; Choudhary & Pucker, 2024). The generated count tables are available via bonndata (Choudhary *et al*., 2025).

Because expression values must be comparable across species and pathways, we implemented a reference-gene-based normalisation strategy:

1. Identification of candidate reference genes: Multiple reference genes exhibiting stable gene expression in *Arabidopsis thaliana* were curated from the literature (**Table S1**).
2. Ortholog detection: Using the phylogenomics approach described above, we identified orthologs of these reference genes in all candidate species. Reference genes with missing orthologs in any species were excluded.
3. Stability check: For each gene, relative expression was computed separately for each species across samples as

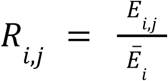

where *E* is the expression of gene *i* in sample *j*, and *E_i_* is the mean expression of that gene across all samples within the same species. Reference genes showing *R_i,j_* ≈1 with low variance across samples were considered stable (**Fig. S19**).
4. Final reference set: Ten genes, which were the most stable across all species (**Table S2**), were selected as internal controls.

For per-sample normalisation, the mean expression of the reference genes (internal controls) in each sample was computed, and all gene counts in that sample were divided by this mean, yielding reference-normalised expression values. Alternative isoforms and close paralogs contributing to the same function were summed to derive total functional gene expression.

Finally, we compared the expression of anthocyanin and proanthocyanidin biosynthesis genes with that of carotenoid biosynthesis genes in both Cucurbitaceae and outgroups to test whether carotenoids might functionally compensate for the absence of anthocyanins in cucurbits.

Mann-Whitney U-test (Mann & Whitney, 1947) was calculated using the implementation in the SciPy library in Python for statistical significance tests. The carotenoid and flavonoid biosynthesis gene candidates used for expression analysis are listed in **Table S3** and **Table S4**, respectively.

## Results

### A highly supported nuclear phylogeny of Cucurbitaceae

In this study, a total of 258 datasets of Cucurbitaceae species were analysed, representing all 15 tribes and 62 genera (out of an estimated 95-115 genera; (Schaefer *et al*., 2008; Schaefer & Renner, 2011b)). An additional 24 outgroup species were included, eight each from Cucurbitales (excluding Cucurbitaceae), Fagales, and Rosales, reflecting their phylogenetic closeness to Cucurbitaceae (The Angiosperm Phylogeny Group *et al*., 2016). Among the Cucurbitaceae datasets, 136 were genomic from various sources, and the remaining were transcriptomic datasets from a previous study (Guo *et al*., 2020). Of the genomic datasets, 82 were newly annotated in this study.

Given the known risk of contamination in all transcriptome assemblies, all 258 Cucurbitaceae datasets were screened for non-cucurbit sequences. Twenty-seven contaminated datasets were identified and excluded from downstream analysis (Choudhary *et al*., 2025)see methods). Using the remaining 255 datasets (including 24 outgroup datasets and multiple accessions per species), we extracted 2805 single-copy orthologs from the eudicotyledons_odb12 BUSCO dataset as phylogenetic markers.

An initial coalescent-based species phylogeny was constructed using FastTree and wASTRAL (**Fig. S1**). Based on this preliminary tree, five datasets were excluded due to low completeness or phylogenetic incongruence: *Xerosicyos perrieri*1 and *Ampelosycios leandrii* (0.1 % and 0% BUSCO, respectively) lacked sufficient gene recovery; *Neoalsomitra pilosa* (9% BUSCO) and *Momordica subangulata* (96.4% BUSCO) clustered in unexpected tribes, likely due to low completeness or misidentified source material; and *Diplocyclos schliebenii* unexpectedly grouped with *Peponium*; whereas *Diplocyclos palmatus* and related taxa clustered with *Coccinia* both in our study and previous studies (Chomicki *et al*., 2020), suggesting possible misidentification or sample contamination of the *D. schliebenii* dataset.

To reconstruct a robust species phylogeny, we adopted a two-method-based approach. First, we generated 2805 gene trees using IQ-TREE under maximum-likelihood (ML) and inferred the species tree using the coalescent method implemented in weighted ASTRAL (wASTRAL) (**Fig. 2**; **Fig. S2**). Second, we performed concatenated ML analyses using a 516-gene supermatrix (**Fig. S3**), as the concatenation method with a relatively small number of loci can reduce systemic errors, whereas very large supermatrices may yield spuriously high support for incorrect topologies (Seo, 2008). Both methods were evaluated for congruence.

**Fig. 2:**
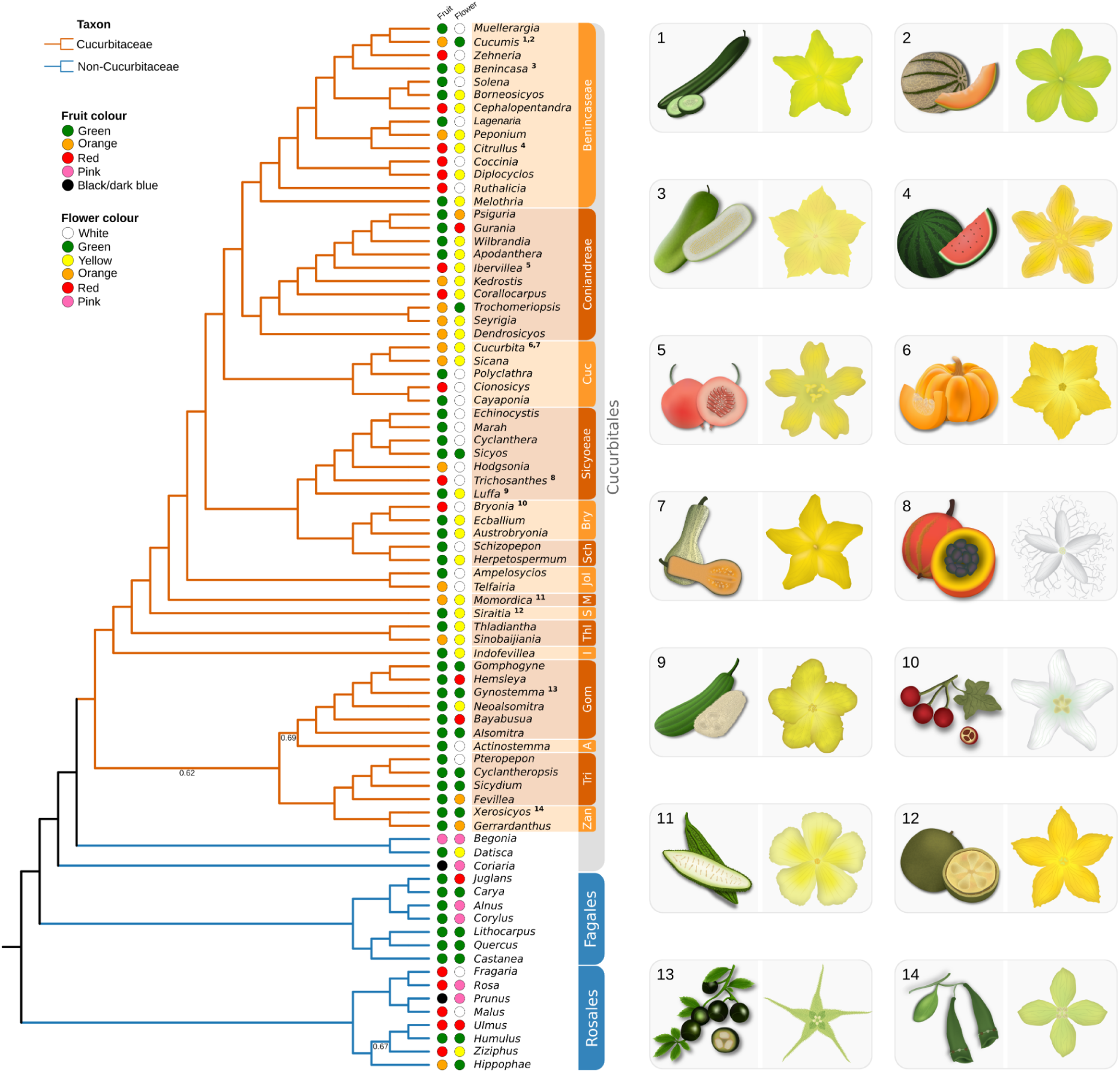
A summary of phylogenetic relationships and pigmentation traits in Cucurbitaceae. Species phylogeny inferred from 2805 BUCO single copy loci, aggregated at the genus level, and representing all 15 tribes of the Cucurbitaceae family. Numbers on branches represent local posterior probability if below 1. Branches corresponding to Cucurbitaceae are shown in orange, while non-Cucurbitaceae outgroups are shown in blue. Fruit and flower colours are mapped onto the terminal tips at the genus level. When multiple colours were observed within a genus, the most intense colour (e.g., red over orange, orange over yellow) was selected. For fruit colour, preference is given to the mature fruit stage and the cut-open fruit section when available. Full species phylogenies are provided in the **Fig. S1, S2 and S3**. Tribe abbreviations: Zan, Zanonieae; Tri, Triceratieae; Act, Actinostemmateae; Gom, Gomphogyneae; I, Indofevilleae; T, Thladiantheae; S, Siraitieae; M, Momordiceae; Jol, Joliffieae; Sch, Schizopeponeae; Bry, Bryonieae; Cuc, Cucurbiteae. On the right, representative Cucurbitaceae species included in this study, illustrating variations in fruits and flowers: 1 *Cucumis sativus*, 2 *Cucumis melo*, 3 *Benincasa hispida*, 4 *Citrullus lanatus*, 5 *Ibervillea tenuisecta*, 6 *Cucurbita pepo*, 7 *Cucurbita argyrosperma*, 8 *Trichosanthes kirilowii*, 9 *Luffa aegyptiaca*, 10 *Bryonia dioica*, 11 *Momordica charantia*, 12 *Siraitia grosvenorii*, 13 *Gynostemma pentaphyllum*, and 14 *Xerosicyos perrieri*.

The coalescent-based tree (**Fig. S2**) provided a more resolved topology and stronger support for monophyletic tribes compared to the concatenated analysis. Branch support was evaluated using local posterior probability (LPP) values. Cucurbitaceae was recovered as a strongly supported monophyletic clade, subdivided into eight major clades (Clade I to Clade VIII). Clade I consists of early diverging lineages Triceratieae, Zanonieae, Actinostemmateae, and Gomphogyneae (LPP=0.62). Clade II consists of Indofevilleae (LPP=1), Clade III-VI containing individual tribes Thladiantheae, Siraitieae, Momordiceae, and Jollifieae (all LPP=1), respectively. Clade VII consists of sister tribes Bryonieae and Scizopeponeae, clustered together as sister to Sicyoeae (LPP=1). Lastly, clade VIII consists of sister tribes Benincaseae and Coniandreae, with Cucurbiteae sister to this group (LPP=1).

The concatenation-based tree largely mirrored this topology, except that the early-diverging lineages formed three individual clades rather than a single combined clade: Triceratie and Zanonieae (Clade I), Gomphogyneae (Clade II), and Actinostemmateae (Clade III). However, the concatenated tree failed to recover monophyly for the *Bryonieae* tribe and has nested phylogenies for *Trichosanthes/Hodgsonia/Cyclanthera* and *Coccinia/Diplocyclos* genera. Therefore, the coalescent-based phylogeny was selected as the most reliable species tree and used for all downstream analyses (**Fig. 2**).

Cucurbitaceae comprise a wide diversity of fruit and vegetable crops (**Fig. 2**). Mapping of fruit and flower pigmentation traits onto the phylogeny (**Fig. 2**) revealed that green, orange and red were the most frequent fruit colours, while white, yellow, and green were predominant among flowers in Cucurbitaceae.

### Absence of major anthocyanin and proanthocyanidin biosynthesis genes in the Cucurbitaceae

Orthologous genes typically retain conserved functions across species and are instrumental for predicting gene functions (Eisen, 1998; Pucker, 2024). Conversely, the inability to identify orthologs in certain lineages can signal lineage-specific gene loss or gene gain in others. To examine this in the context of pigmentation in Cucurbitaceae, we performed a comprehensive ortholog-based survey of flavonoid biosynthesis genes (FBGs) across all Cucurbitaceae datasets included in this study. The analysis incorporated both sequence similarity to functionally characterised reference genes and conservation of key amino acid residues essential for enzymatic activity. In addition, phylogenetic analyses were performed for each major structural and regulatory gene family to validate orthology and infer Cucurbitaceae pigmentation traits.

Consistent with expectations, the core flavonoid and phenylpropanoid pathway genes (e.g., *PAL*, *C4H*, *4CL*, *CHS*, *CHI*, *FNS II*, *F3H*, *F3’H*, and *FLS*) were readily identified across all Cucurbitaceae datasets, displaying clear orthology with known functional genes in other eudicots. However, the analyses revealed a striking absence of key anthocyanin and proanthocyanidin biosynthetic genes, *DFR*, *ANS*, and *arGST* for anthocyanin biosynthesis and *ANR* and *LAR* for proanthocyanidin biosynthesis (**Fig. 3a-j, Fig. S4-S13**). A few exceptions were observed among the early diverging species in the Cucurbitaceae, where *ANR* was detected in *Bayabusua clarkie*, *LAR* was detected in *Alsomitra macrocarpa* and *B. clarkie*, and *DFR* or DFR-like sequences in *B. clarkie*, *A. macrocarpa*, *Gynostemma* and a few related taxa. In contrast, the vast majority of Cucurbitaceae lacked all essential structural genes required for anthocyanidin and proanthocyanidin biosynthesis, suggesting a family-wide irreversible loss of both branches of the flavonoid pathway. The status of *U3GT* (UDP-glucose: flavonoid 3-O-glucosyltransferase) could not be conclusively resolved. Although Cucurbitaceae sequences clustering with canonical *U3GT* gene sequences were identified (**Fig. 3h**), some glycosyltransferases are known to exhibit substrate promiscuity, often catalysing the glycosylation of both flavonols and anthocyanidins (Lee *et al*., 2005). The functional role of *U3GT*-like sequences remains uncertain in Cucurbitaceae, requiring enzymatic validation. To investigate the regulatory control, the MBW complex was examined. Orthologs of subgroup 6 (SG6) MYBs, which regulate anthocyanin biosynthesis, and PA1 MYB and PA2 MYB (SG5), controlling proanthocyanidin biosynthesis, were entirely absent from Cucurbitaceae. Subgroup 7 (SG7) MYBs, which regulate flavonol biosynthesis, were present and phylogenetically conserved. The other two components of the MBW complex, bHLH and WD40 proteins, were detected in almost all datasets, consistent with their pleiotropic roles in diverse biological processes beyond flavonoid biosynthesis.

**Fig. 3:**
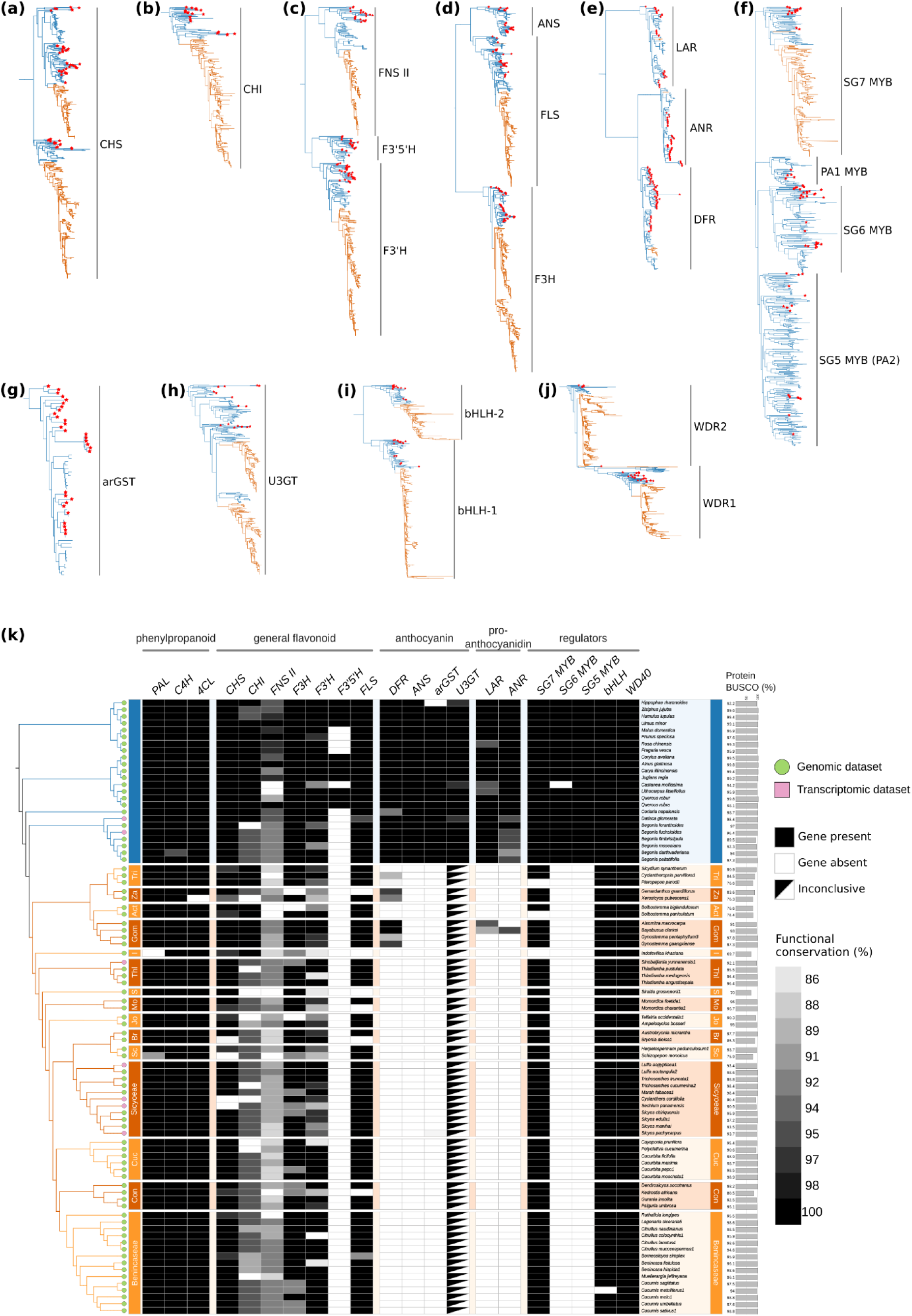
The phylogenetic pattern of major flavonoid pathway genes in Cucurbitaceae. Maximum-likelihood phylogenetic trees of major flavonoid pathway genes are shown at the top. Branches corresponding to Cucurbitaceae sequences are shown in orange, while non-Cucurbitaceae outgroups are shown in blue. Red stars at the tip of terminal branches indicate functional enzymatic sequences identified in previous studies (Choudhary *et al*., 2025). Gene trees for (a) CHS, (b) CHI, (c) CYP450 family (FNS II, F3’H, and F3’5’H), (d) 2-ODD family (F3H, FLS, and ANS), (e) SDR family (LAR, ANR, and DFR), (f) flavonoid-related MYBs (SG7 MYB, SG6 MYB, SG5 MYB, and PA1 MYB), (g) arGST, (h) U3GT, (i) flavonoid-related bHLHs, and (i) flavonoid-related WD40 are shown. Full phylogenetic trees are provided in **Fig. S4-13**. On the bottom, the presence-absence summary matrix of flavonoid biosynthesis genes in some representative Cucurbitaceae species mapped to the Cucurbitaceae phylogeny is shown (k). Outgroups (blue) and Cucurbitaceae (orange) species are annotated with genomic (green tips) and transcriptomic (pink tips) data sources. Gene presence, absence, or putative non-functional sequences are shown as black, white and grey boxes, respectively. The rightmost bars represent the percentage of conserved BUSCO genes identified in the polypeptide sequences of each dataset shown. Tribe abbreviations: Tri, Triceratieae; Za, Zanonieae; Act, Actinostemmateae; Gom, Gomphogyneae; I, Indofevilleae; Thl, Thladiantheae; S, Siraitieae; Mo, Momordiceae; Jo, Joliffieae; Br, Bryonieae; Sc, Schizopeponeae; Cuc, Cucurbiteae; Con, Coniandreae. For the full presence-absence matrix across all datasets analysed in this study, see **Fig. S14**.

The identified presence-absence patterns were mapped onto the Cucurbitaceae species phylogeny (**Fig. 3k**). While interpreting gene absence as evolutionary loss is vulnerable to technical artefacts like incomplete assemblies, low sequence coverage or extreme sequence divergence, several lines of evidence support the inferred evolutionary gene loss. First, consistent absence across multiple independent, high-quality genome assemblies and annotations; second, concordant absence in both genomic and transcriptomic datasets from diverse lineages; and lastly, the identification of *DFR*, *LAR*, and *ANR* in a few early-diverging taxa using the same approach.

### Microsynteny supports the loss of anthocyanin biosynthesis genes

The evolutionary path of anthocyanin and proanthocyanidin genes might help in understanding the inability to find anthocyanidin and proanthocyanidin biosynthesis genes in Cucurbitaceae. To elucidate the basis for the absence of anthocyanin and proanthocyanidin biosynthesis genes in Cucurbitaceae, we conducted a comparative microsynteny analysis focused on key anthocyanin biosynthetic loci.

Before examining fine-scale microsynteny, it was necessary to understand the large-scale genomic context, particularly the history of whole-genome duplication (WGD) events in the analysed species relative to the reference genome (*Fragaria vesca*). All eudicots share an ancient core eudicot common hexaploidization (ECH) event, estimated to have occurred approximately 115-130 million years ago (Mya) (Jaillon *et al*., 2007). Following this, the Cucurbitales common tetraploidization (CCT) event, shared among all members of the order, occurred around 93-105 Mya (Wang *et al*., 2022). Begoniaceae experienced an additional WGD around 35 Mya (Li *et al*., 2022). Within Cucurbitaceae, additional lineage-specific WGDs have been reported. A major event, the *Cucurbita*-specific tetraploidization (CST), occurred around 30 Mya, resulting from an ancient allotetraploidization followed by extensive gene loss and rediploidization (Montero-Pau *et al*., 2018). Furthermore, a distinct Sicyoeae-specific WGD, dated to approximately 25 Mya (Fu *et al*., 2021), was previously identified and is shared among *Sicyos* and closely related taxa (Guo *et al*., 2020).

Given this complex history of polyploidization, accurately inferring orthology required representing one-to-many syntenic relationships in comparisons with the *F. vesca* reference genome sequence. Synteny depth analyses confirmed the expected orthology ratios that reflect these duplication events: 1:1 between *F. vesca* and *Quercus robur*, 1:2 between *F. vesca* and most Cucurbitaceae; and 1:4 between *F. vesca* and the highly polyploid lineages *Begonia*, *Cucurbita*, and *Sicyos* (**Fig. S16**). Unexpectedly, we observed a 1:4 orthology ratio between *F. vesca* and *C. nepalensis*. Because *Coriaria* is known to share only the CCT event, a 1:2 ratio would have been expected. This pattern instead suggests an additional, lineage-specific WGD in *Coriaria*. We tested this hypothesis and indeed detected a previously suggested (Zhao *et al*., 2023), independent whole-genome duplication event, which we dated at 18 Mya, providing strong support for *Coriaria*-specific WGD (**Fig. S17**).

With the polyploidization history clarified, we next investigated the conservation of genomic regions harbouring key anthocyanin biosynthesis genes. Comparative analyses across 19 high-quality genome sequences (including both outgroups and Cucurbitaceae) revealed well-conserved syntenic blocks surrounding the *DFR*, *ANS*, and *arGST* loci (**Fig. 4**), as well as for proanthocyanidin biosynthesis genes *ANR* and *LAR* (**Fig. S18**). To validate the reliability of the identified syntenic relationships, we also generated gene trees to confirm identified homologs represented true orthologous relationships.

**Fig. 4:**
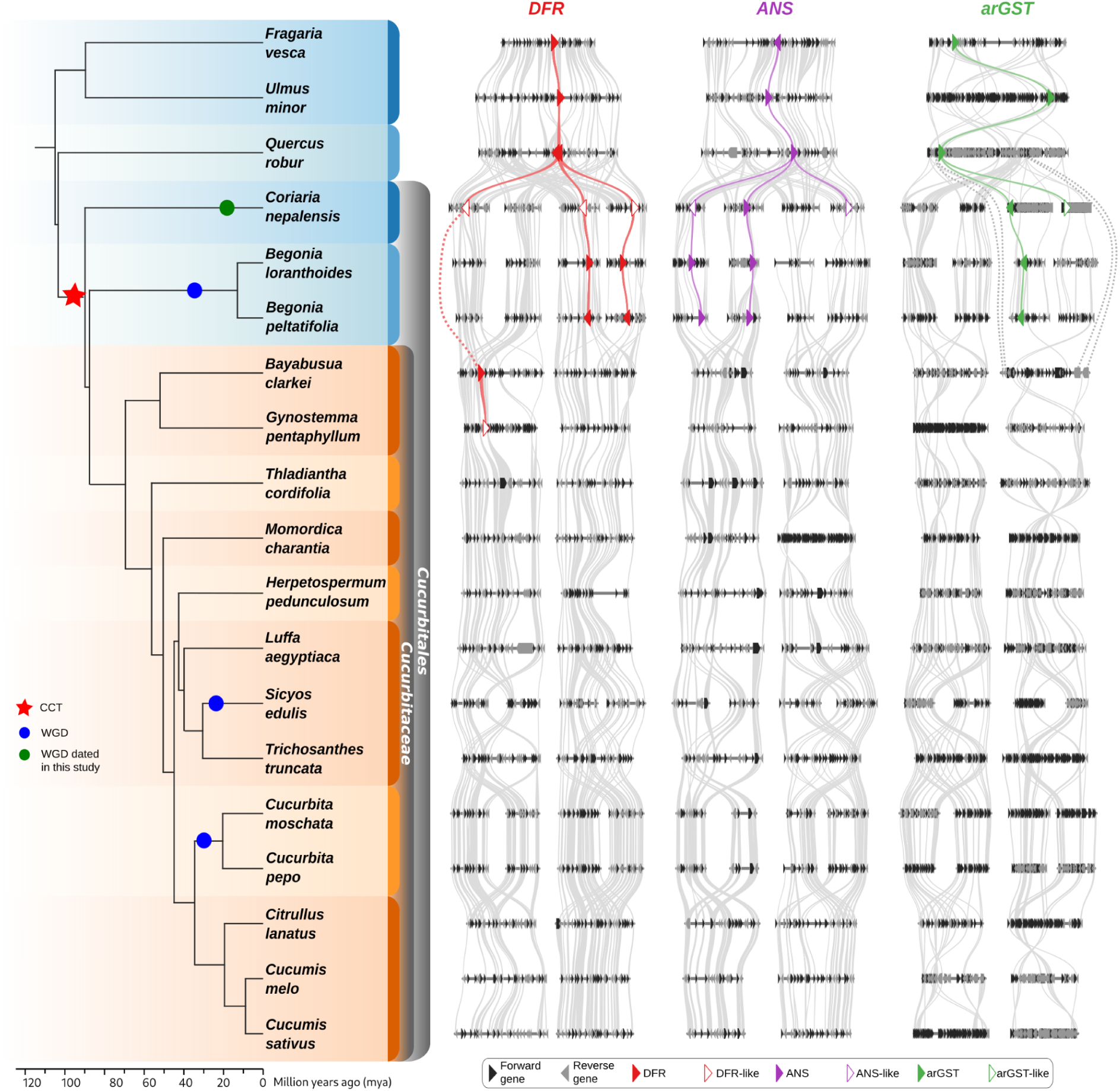
Microsynteny around DFR, ANS, and arGST loci. On the left, the species phylogeny is shown. A red star denotes the Cucurbitales common tetraploidization (CCT) event, while blue circles indicate lineage-specific whole-genome duplications (WGDs): the Begoniaceae common tetraploidization (BCT), the Sicyoeae WGD, and the Cucurbita-specific polyploidization (CST). A green circle marks the putative Coriaria-specific WGD identified in this study. On the right, conserved syntenic regions for the anthocyanin biosynthesis genes, *DFR* (red), *ANS* (purple), and *arGST* (green) are shown. Genes outlined with unfilled coloured borders represent putatively non-functional copies, based on conserved amino acid residue analysis. Other genes are shown in black (forward strand) and grey (reverse strand). The number of syntenic blocks displayed per species reflects the degree of collinearity relative to *F. vesca* and corresponds to the number of polyploidization events in the evolutionary history of each lineage.

The microsynteny pattern clearly supports a loss of the anthocyanin biosynthesis genes *DFR*, *ANS*, and *arGST* across Cucurbitaceae. A *DFR* homolog was retained in *Bayabusua clarkie,* and a DFR-like sequence was identified in *Gynostemma pentaphyllum* (early diverging Cucurbitaceae). Similarly, the proanthocyanidin genes *LAR* and *ANR* were absent from all Cucurbitaceae examined except for *LAR* in a few early-diverging taxa. This suggests that the anthocyanin biosynthesis gene loss occurred at the base of the Cucurbitaceae family, while the proanthocyanidin gene loss occurred after the divergence of basal cucurbit lineages.

Together, the phylogenetic trees, biosynthetic gene identification, and microsynteny provide robust genomic support for the complete loss of both anthocyanidin and proanthocyanidin biosynthetic genes in the majority of Cucurbitaceae.

### Carotenoids as functional replacements for anthocyanins

The complete loss of the anthocyanin and proanthocyanidin biosynthetic genes in Cucurbitaceae raises two key questions: first, ‘what is the evolutionary fate of the upstream flavonoid biosynthetic pathway?’ and second, ‘have carotenoids, the other major class of plant pigments, functionally compensated for anthocyanin loss?’.

To assess whether the upstream flavonoid genes remain transcriptionally active, we performed comparative expression analysis using RNA-seq data from eight representative Cucurbitaceae species and eight phylogenetically close outgroup species from *Fagales* and *Rosales*. Across all Cucurbitaceae species, we observed a marked reduction in transcript abundance for the remaining flavonoid biosynthesis genes (**Fig. 5a**). Particularly, *CHS*, which encodes the enzyme catalysing the first committed step of flavonoid biosynthesis, exhibited near-negligible expression levels in Cucurbitaceae species, suggesting that the entire flux into the flavonoid pathway is effectively constrained at its entry point. While *CHI*, *F3’H* and *FLS* retained residual expression, their overall transcript levels were significantly lower (p<0.001) in Cucurbitaceae than those observed in taxa with functional anthocyanin biosynthesis (**Fig. 5b**). These results might imply that the upstream flavonoid biosynthesis genes in Cucurbitaceae are undergoing relaxed or negative purifying selection, consistent with functional redundancy, and gradual pathway erosion following the loss of anthocyanin and proanthocyanidin branches.

**Fig. 5:**
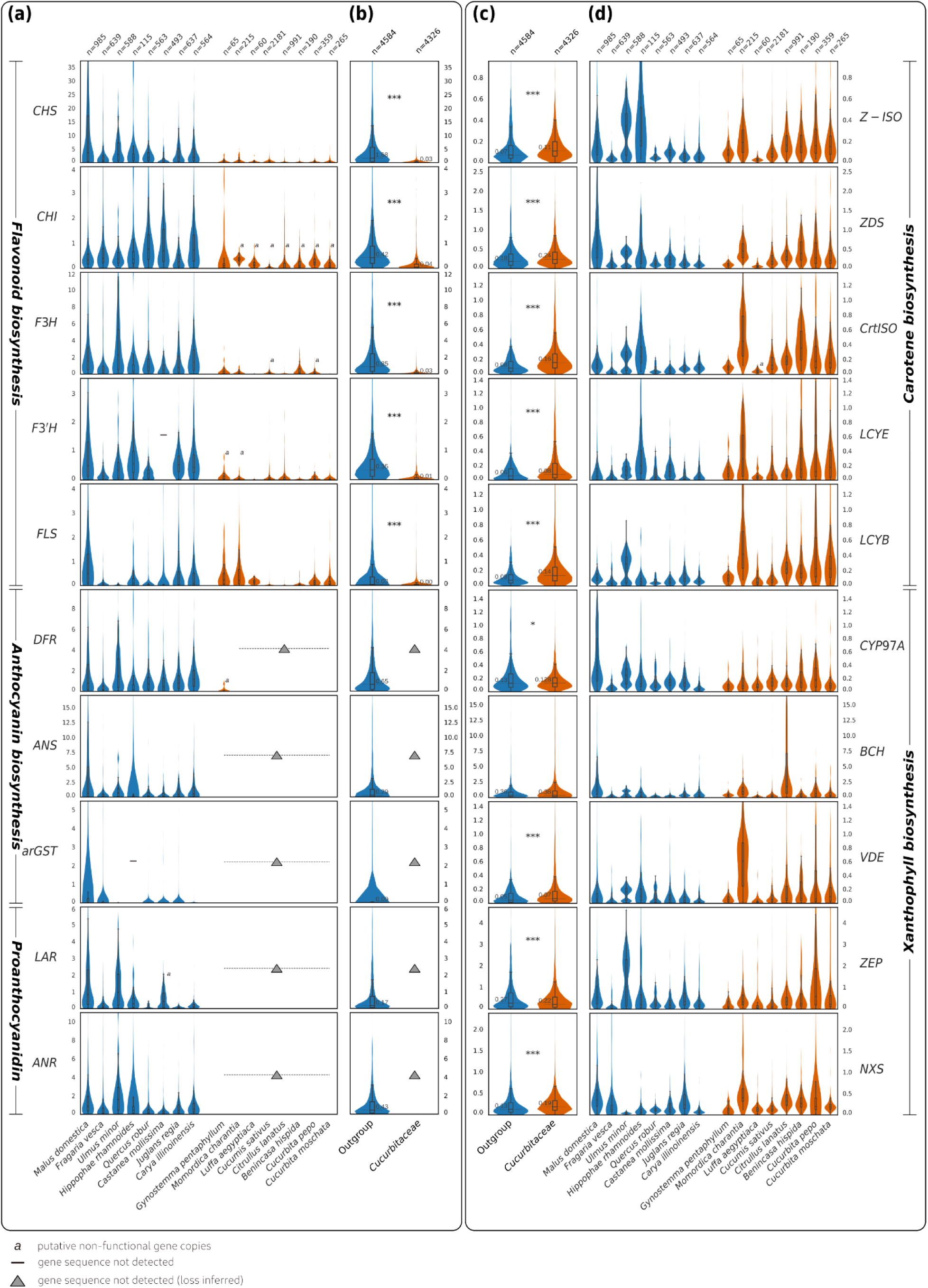
(a) Expression analysis of core flavonoid biosynthetic genes in Cucurbitaceae (orange) and outgroups (blue). Sample size (*n*) for each group is indicated above the plots. Superscript ‘*a’* next to specific violin plots denote putative non-functional gene copies inferred from conserved amino acid residue analyses (**Table S3** and **S4**). Grey triangles indicate gene sequence not detected (loss inferred), while a black dash represents gene sequence not detected. (b) Combined expression analysis of flavonoid biosynthesis genes between all Cucurbitaceae and outgroups. (c) Combined expression analysis of carotenoid biosynthesis genes between all Cucurbitaceae and outgroups. Median expression value for both groups (Cucurbitaceae and outgroup) is denoted on the plot. Statistical significance is denoted as ‘***’ (p <=0.001),‘**’ (p<=0.01), and ‘*’ (p<=0.05). (d) Expression analysis of the core carotenoid biosynthetic genes in outgroups (blue) and Cucurbitaceae (orange). Boxplots within violin plots indicate the median, interquartile range, and outliers.

Given the multifunctional roles of anthocyanins in pigmentation, photoprotection, and pollinator attraction, we next investigated whether carotenoids may have functionally compensated for the anthocyanin loss. To test this hypothesis, we compared the expression of core carotenoid biosynthetic genes between Cucurbitaceae and the same outgroup taxa (**Fig. 5d**). Although carotenoid biosynthesis represents a primary metabolic pathway that is constitutively active in photosynthetic tissues, Cucurbitaceae exhibited a slightly higher overall transcript abundance across most carotenoid biosynthetic genes compared to outgroups (p < 0.01; **Fig. 5c**). This elevated expression may suggest adaptive enhancement of carotenoid production, both to compensate for the visual signalling roles and to maintain photoprotection in the absence of anthocyanins and possibly most other flavonoids. The observed transcript abundance is consistent with the high carotenoid content reported in cucurbit fruits and flowers.

## Discussion

Gene loss is a major evolutionary force shaping morphological diversity, ecological adaptation, and lineage-specific innovation across organisms (Smith & Rausher, 2011; Albalat & Cañestro, 2016). Although often associated with pseudogenization and loss of function, gene loss can also drive evolutionary innovation, where the removal of redundant or dispensable genes allows reconfiguration of metabolic or developmental networks, promoting novel adaptive functions (Olson, 1999; Sharma *et al*., 2018).

In this study, we report an exceptional case of large-scale pathway gene loss defining the pigmentation history of the Cucurbitaceae family. Our analysis reveals the almost complete absence of core structural genes required for anthocyanin and proanthocyanidin biosynthesis, together with the loss of regulatory SG6 and SG5 MYB transcription factors responsible for activating these pathways. Such a comprehensive loss of both structural and regulatory components across an entire plant family is exceedingly rare and represents a fundamental evolutionary shift in the pigmentation mechanism in Cucurbitaceae.

Anthocyanins play central ecological roles by producing diverse colours that attract pollinators and seed dispersers, while proanthocyanidins contribute to defence, seed coat pigmentation, and dormancy (Dixon *et al*., 2005; Grünig *et al*., 2025). Given that many extant species remain insect- and bird-pollinated (Schaefer & Renner, 2011a), the complete absence of these pathways is remarkable. However, several early-diverging Cucurbitaceae lineages, such as *Bayabusua*, *Alsomitra*, *Pteropepon*, and *Siolmatra*, are wind-pollinated, suggesting reduced reliance on visual signals for pollination and seed dispersal. This shift in pollination strategy may have rendered anthocyanin pigmentation functionally irrelevant early in Cucurbit evolution, facilitating relaxed selection pressure and eventual gene loss.

Gene loss is an irreversible evolutionary process that prevents reacquisition of a trait through the same genetic basis, thus opening possibilities for compensatory or convergent evolutionary solutions (Pucker *et al*., 2024; Khatun *et al*., 2025). Following the initial loss of anthocyanin-based pigmentation, Cucurbitaceae appear to have evolved alternative strategies for pollination and seed dispersal. The family exhibits remarkable evolutionary innovation in seed dispersal strategies. Autochory, the self-dispersal of seeds with gravity or ballistic movements, has been reported in some cucurbits. Species such as *Alsomitra macrocarpa* and *Bayabusua clarkei* produce wing-shaped seeds capable of gliding hundreds of meters (Certini, 2023). In *Ecballium elaterium* (squirting cucumber), fruit ripening is followed soon by ballistic ejection of seeds reaching several metres far (Box *et al*., 2024; Gorges *et al*., 2025). In wild cucurbits, endozoochory, i.e, ingestion of fruits by animals and defecation of the seeds elsewhere, has been proposed as a major dispersal route, with ancestral species uniquely adapted to dispersal by large herbivores (Kistler *et al*., 2015; Johnson *et al*., 2025). In addition, several genera, including *Luffa*, *Cayaponia*, *Hodsgonia*, and *Sicyos,* are adapted to water dispersal (Ridley, 1930). These seed-dispersal innovations may have been associated with reduced visual signals, resulting from anthocyanin loss.

Carotenoids, which produce yellow, red, and orange pigmentation in Cucurbitaceae fruits and flowers, likely assumed the visual signalling functions of anthocyanins. The predominance of carotenoid-based colouration across domesticated Cucurbitaceae, combined with elevated carotenoid biosynthetic gene expression, supports this hypothesis. Similarly, morphological innovations such as fringed petal lobes in genera like *Trichosanthes* (de Boer *et al*., 2012) may have evolved to enhance pollinator attraction through visual or tactile cues, partially compensating for the absence of anthocyanin pigments. Beyond visual attraction, seed pigmentation and defence functions previously attributed to proanthocyanidins may have been replaced by other biochemical mechanisms. One potential candidate is melanin, a lesser-studied plant pigment already known to colour the black seeds of *Citrullus lanatus* (Glagoleva *et al*., 2020).

While our findings reveal a clear pattern of gene loss, the temporal and phylogenetic context of these events remains to be fully resolved. Our species phylogeny indicates that the early-diverging tribes, Triceratieae, Zanonieae, Actinostemmateae, and Gomphogyneae, form a monophyletic basal clade (Clade I), consistent with previous reconstructions (Schaefer *et al*., 2008; Lin *et al*., 2025). However, alternative topologies reported for these early-diverging taxa (Chomicki *et al*., 2020; Guo *et al*., 2020) highlight the need for denser taxon sampling and higher-quality genome sequences. Improved genomic resolution in these early lineages will be essential for pinpointing the exact timing and order of gene loss events.

The detection of *DFR*, *LAR*, and *ANR* orthologs in *Bayabusua* and related early-diverging taxa suggests that the proanthocyanidin branch may have persisted longer than the anthocyanin branch. This pattern supports a stepwise loss, in which anthocyanin biosynthesis was lost first, potentially following the loss of its specific regulatory MYB activators, while the proanthocyanidin pathway remained partially functional before being completely lost in later-diverging Cucurbitaceae. This hypothesis aligns with previous studies identifying loss of regulatory MYBs as the most observed causal factor behind flavonoid pathway blocks (Wheeler *et al*., 2022; Marin-Recinos & Pucker, 2024). Testing this model will require multiple high-quality genome sequences from early-branching lineages to confirm whether the remaining flavonoid biosynthesis genes are functional or represent degenerated relics.

The evolutionary loss of anthocyanin genes in Cucurbitaceae thus provides a unique comparative framework for identifying novel anthocyanin-biosynthesis genes by contrasting Cucurbitaceae (anthocyanin-deficient) and anthocyanin-producing lineages. Thereby, we propose Cucurbitaceae as an exceptional model system for studying the evolution, regulation, and diversification of anthocyanin and proanthocyanidin biosynthesis - the colourful branches of the flavonoid pathway.

In summary, our findings reveal a systematic loss of anthocyanin and proanthocyanidin genes in Cucurbitaceae and indicate a compensatory enhancement of the carotenoid pigmentation and alternative ecological strategies. By uncovering the genomic and phylogenetic foundation of this transition, our study provides a framework for understanding the evolution of pigmentation systems in plants and challenges the long-held assumption that anthocyanin pigmentation is (universally) conserved across flowering plants.

## Acknowledgements

This work was supported by the de.NBI Cloud within the German Network for Bioinformatics Infrastructure (de.NBI) and ELIXIR-DE (Forschungszentrum Jülich and W-de.NBI-001, W-de.NBI-004, W-de.NBI-008, W-de.NBI-010, W-de.NBI-013, W-de.NBI-014, W-de.NBI-016, W-de.NBI-022). We thank all members of the research group Plant Biotechnology and Bioinformatics for their discussion and support. We acknowledge support from Project DEAL and the University of Bonn for open access publication.

## Author contributions

BP and NC designed the project. NC conducted the analyses and BP supervised the work. MH designed illustrations. NC wrote the manuscript. NC, MH, and BP revised the manuscript, and all authors agreed to its submission.

## Data availability

All data sets underlying this study are publicly available. The data supporting the results and conclusions are included in the article and its supporting information. All sequences and additional dataset files are accessible via bonndata (https://doi.org/10.60507/FK2/ZNVSLA). Customised Python scripts for the analyses in this study are available through GitHub (https://github.com/NancyChoudhary28/Cucurbits).

## Supporting information

### Description

**Fig. S1** Coalescence species tree constructed from 2805 BUSCO orthogroups across 255 non-contaminated datasets.

**Fig. S2** Coalescence species tree constructed from 2805 BUSCO orthogroups across 250 non-contaminated datasets.

**Fig. S3** The ML tree reconstructed from a concatenated supermatrix of 516 BUSCO orthogroups across 250 non-contaminated datasets.

**Fig. S4** Codon-based maximum-likelihood phylogenetic tree of CHS sequences constructed with **(a)** MAFFT and IQ-TREE, and **(b)** MUSCLE and IQ-TREE

**Fig. S5** Codon-based maximum-likelihood phylogenetic tree of CHI sequences constructed with **(a)** MAFFT and IQ-TREE, and **(b)** MUSCLE and IQ-TREE

**Fig. S6** Codon-based maximum-likelihood phylogenetic tree of CYP450 sequences constructed with **(a)** MAFFT and IQ-TREE, and **(b)** MUSCLE and IQ-TREE

**Fig. S7** Codon-based maximum-likelihood phylogenetic tree of 2ODD sequences constructed with **(a)** MAFFT and IQ-TREE, and **(b)** MUSCLE and IQ-TREE

**Fig. S8** Codon-based maximum-likelihood phylogenetic tree of SDR sequences constructed with **(a)** MAFFT and IQ-TREE, and **(b)** MUSCLE and IQ-TREE

**Fig. S9** Codon-based maximum-likelihood phylogenetic tree of arGST sequences constructed with **(a)** MAFFT and IQ-TREE, and **(b)** MUSCLE and IQ-TREE

**Fig. S10** Codon-based maximum-likelihood phylogenetic tree of U3GT sequences constructed with **(a)** MAFFT and IQ-TREE, and **(b)** MUSCLE and IQ-TREE

**Fig. S11** Codon-based maximum-likelihood phylogenetic tree of MYBs sequences constructed with **(a)** MAFFT and IQ-TREE, and **(b)** MUSCLE and IQ-TREE

**Fig. S12** Codon-based maximum-likelihood phylogenetic tree of bHLH sequences constructed with **(a)** MAFFT and IQ-TREE, and **(b)** MUSCLE and IQ-TREE

**Fig. S13** Codon-based maximum-likelihood phylogenetic tree of WD40 sequences constructed with **(a)** MAFFT and IQ-TREE, and **(b)** MUSCLE and IQ-TREE

**Fig. S14** Presence-Absence Matrix of Flavonoid biosynthesis genes in all datasets used in this study

**Fig. S15** Contamination profiles of *Cucurbita pepo*3 and *Sicyosperma gracile*

**Fig. S16** Synteny depth plots of all reference vs target datasets used in the microsynteny analysis

**Fig. S17** Whole Genome Duplication (WGD) detection and dating in *Coriaria nepalensis*.

**Fig. S18** Microsynteny around *LAR* and *ANR* loci

**Fig. S19** Relative expression stability of selected reference genes across 16 plant species

**Table S1** Candidate reference genes evaluated for expression normalisation

**Table S2** Reference gene orthologs used for expression normalisation across species

**Table S3** Genes used for expression analyses of carotenoid biosynthesis genes in outgroups and Cucurbitaceae

**Table S4** Genes used for expression analyses of flavonoid biosynthesis genes in outgroups and Cucurbitaceae

