## Supplementary material for "Out of the blue: Family-wide loss of anthocyanin biosynthesis in Cucurbitaceae": Fig. S2

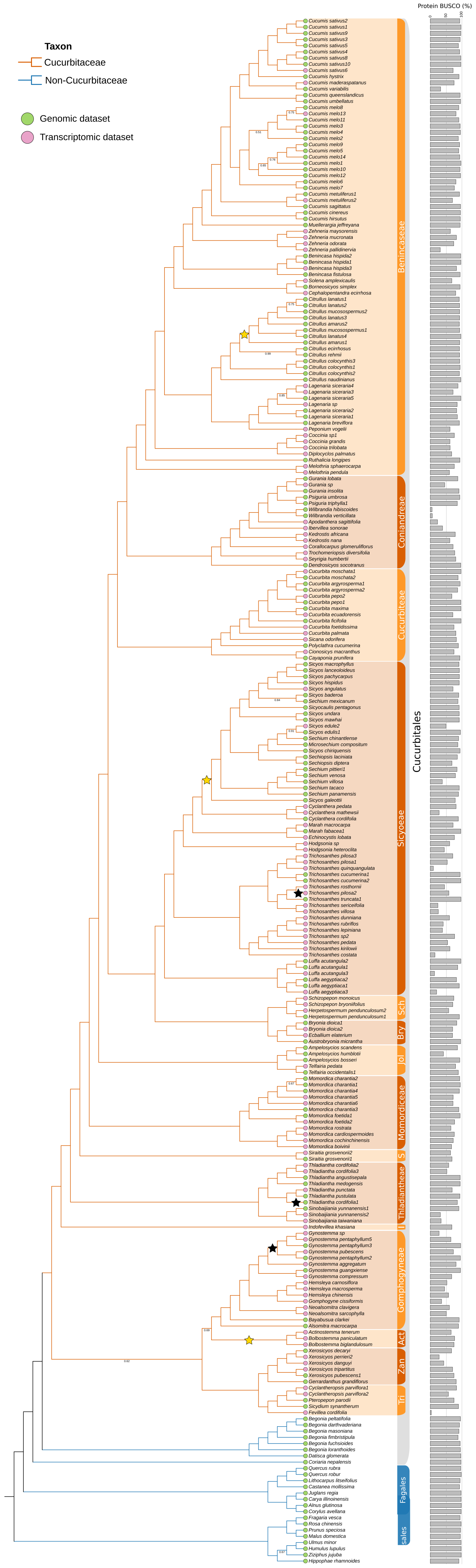


**Fig. S2: Coalescence species tree constructed from 2805 BUSCO orthogroups across 250 non-contaminated datasets.**

Numbers on branches represent local posterior probability if below 1. The 15 Cucurbitaceae tribes are highlighted with orange coloured backgrounds. Branches corresponding to Cucurbitaceae are shown in orange, while non-Cucurbitaceae outgroups are shown in blue. Dataset type (genomic or transcriptomic) is indicated at the terminal tips in green and pink, respectively. The percentage of conserved BUSCO genes identified in the polypeptide sequences of each dataset is shown on the far right. Species names follow Plants of the World Online (POWO; powo.science.kew.org). Species/clades marked with black stars are retained but warrants caution. For example, multiple accessions of *Gynostemma pentaphyllum* did not form a monophyletic clade; instead, *G. pubescens* was nested among them, which may suggest incomplete lineage sorting, introgression between the two species, or potential misidentification. Similary, accession of *Thladiantha cordifolia* (1,2,3) and *Trichosanthes pilosa* (1,2,3) failed to form monophyletic groups. The species in the clade marked with yellow stars were merged into a single genus based on previous phylogenetic positions (Schaefer and Renner, 2011): *Bolbostemma* in *Actinostemma*; *Microsechium, Sicyocaulis, Sicyosperma, Sechiopsis*, and *Sechium* were all subsumed within *Sicyos*. The clustering of *C. lanatus* with *C. mucosospermus* is consistent with previous phylogenetic studies (Chomicki & Renner, 2015). Tribe abbreviations: Act, Actinostemmateae; Tri, Triceratieae; Zan, Zanonieae; I, Indofevilleae; S, Siraitieae; Jol, Joliffieae; Bry, Bryonieae; Sch; Schizopeponeae
