## Supplementary material for "Out of the blue: Family-wide loss of anthocyanin biosynthesis in Cucurbitaceae": Fig. S6b

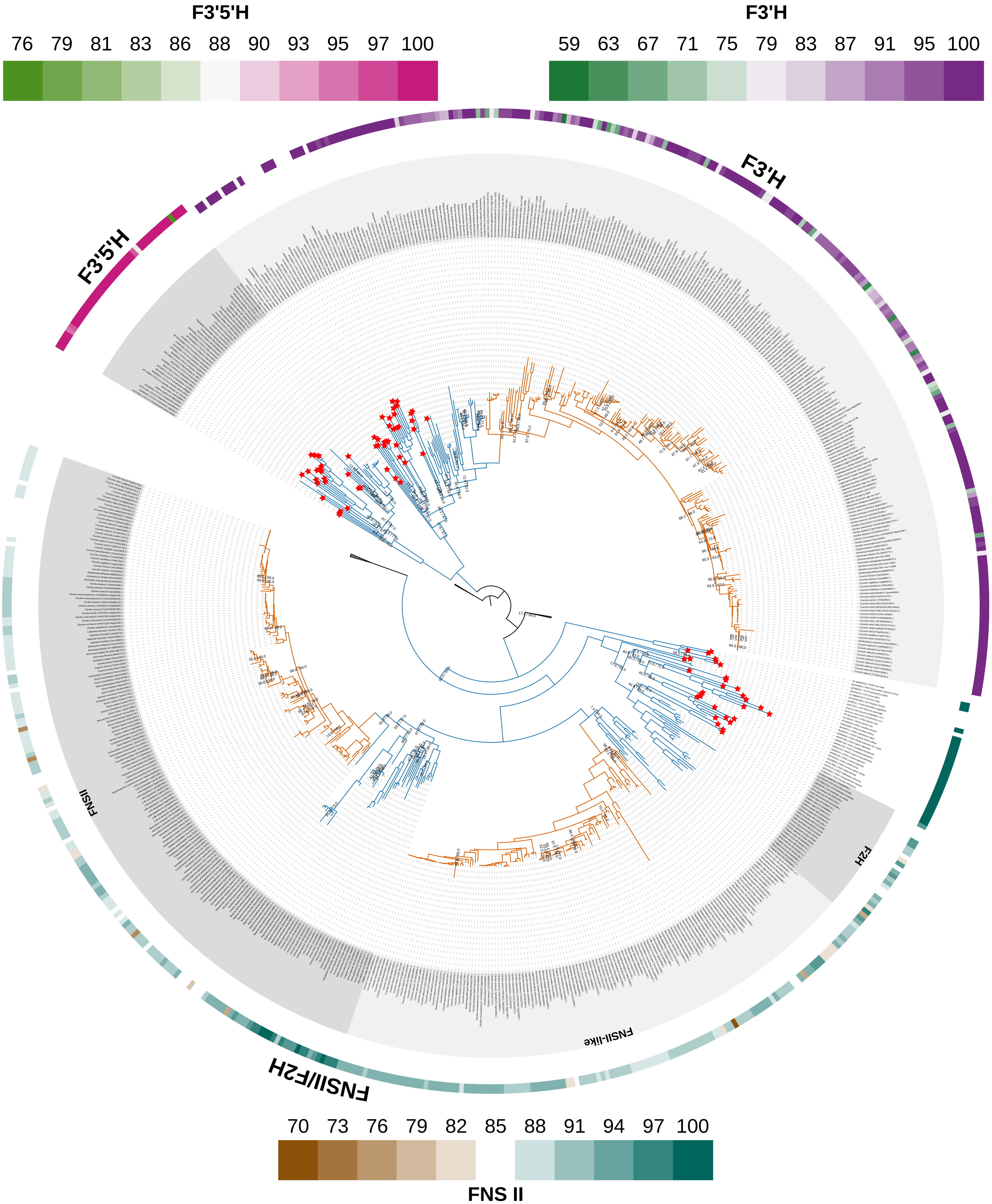


**Fig. S6b: Codon-based maximum-likelihood phylogenetic tree of CYP450 sequences constructed with MUSCLE and IQ-TREE.**

The best-fit substitution model selected by Model-Finder was GTR+F+I+R10. Cucurbitaceae sequences are shown in orange branches, and non-Cucurbitaceae sequences shown in blue. Red stars at the tips of terminal branches indicate functional enzymatic sequences identified in previous studies. Numbers above nodes represent Shimodaira-Hasegawa approximate likelihood ratio test (SH-aLRT) support (%)/ ultrafast bootstrap support (%) based on 1000 replicates. Only percentages below 100 are shown. The outermost colour gradient indicates the percentage of conserved functionally important residues for the corresponding function, thereby reflecting the degree of functional conservation of each sequence. F3'H, flavonoid 3'-hydroxylase; F3'5'H, flavonoid 3',5'-hydroxylase; F2H, flavanone 2-hydroxylase; FNSII, flavone synthase II
