## Supplementary material for "Out of the blue: Family-wide loss of anthocyanin biosynthesis in Cucurbitaceae": Fig. S10a

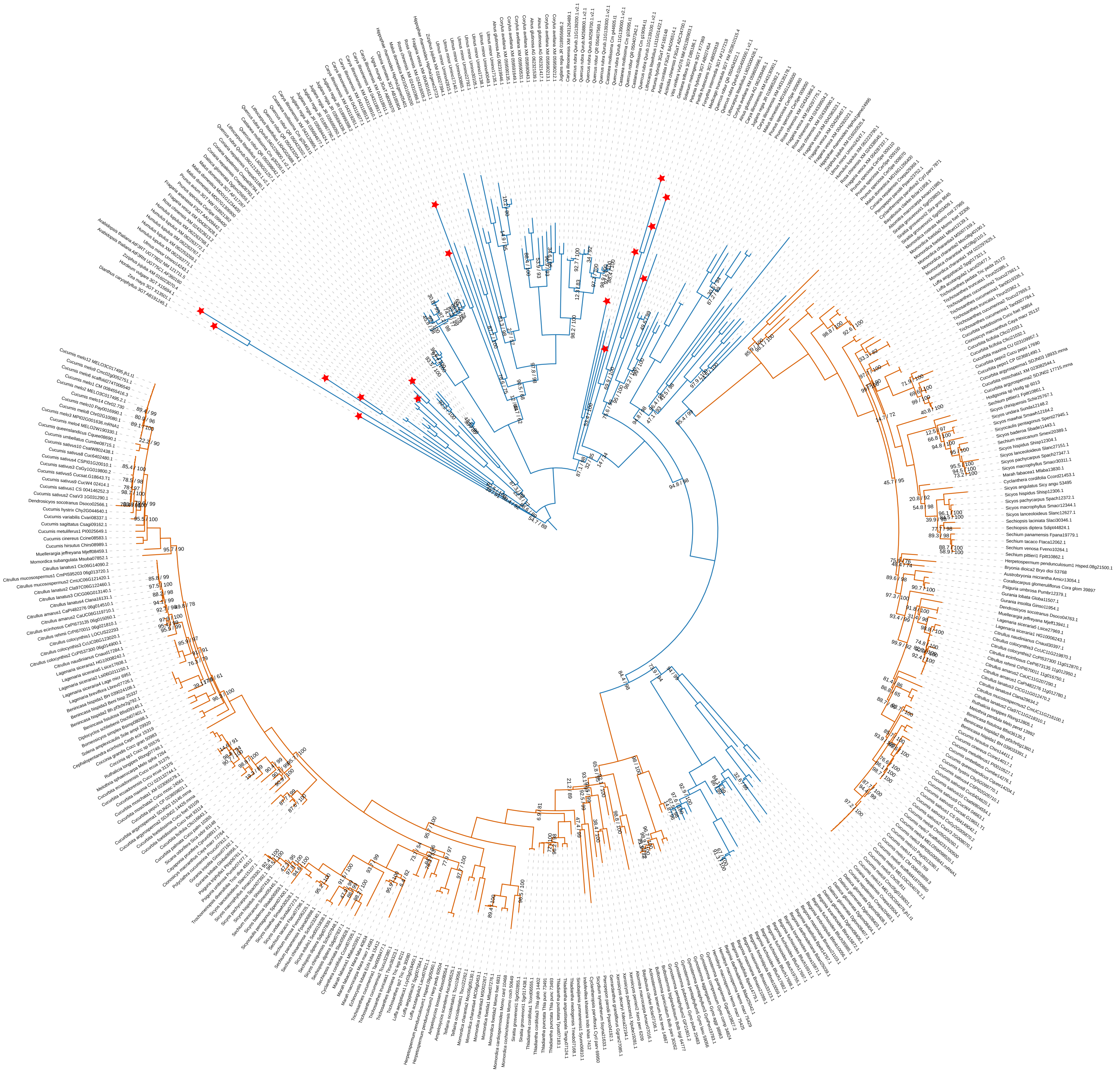

**Fig. S10a: Codon-based maximum-likelihood phylogenetic tree of U3GT sequences constructed with MAFFT and IQ-TREE.**

The best-fit substitution model selected by Model-Finder was TVMe+I+R6. Cucurbitaceae sequences are shown in orange branches, and non-Cucurbitaceae sequences shown in blue. Red stars at the tip of terminal branches indicate functional enzymatic sequences identified in previous studies. Numbers above nodes represent Shimodaira-Hasegawa approximate likelihood ratio test (SH-aLRT) support (%)/ ultrafast bootstrap support (%) based on 1000 replicates. Only percentages below 100 are shown. U3GT, UDP-glucose-dependent 3-*O*-glucosyltransferase
