## Supplementary material for "Out of the blue: Family-wide loss of anthocyanin biosynthesis in Cucurbitaceae": Fig. S11a

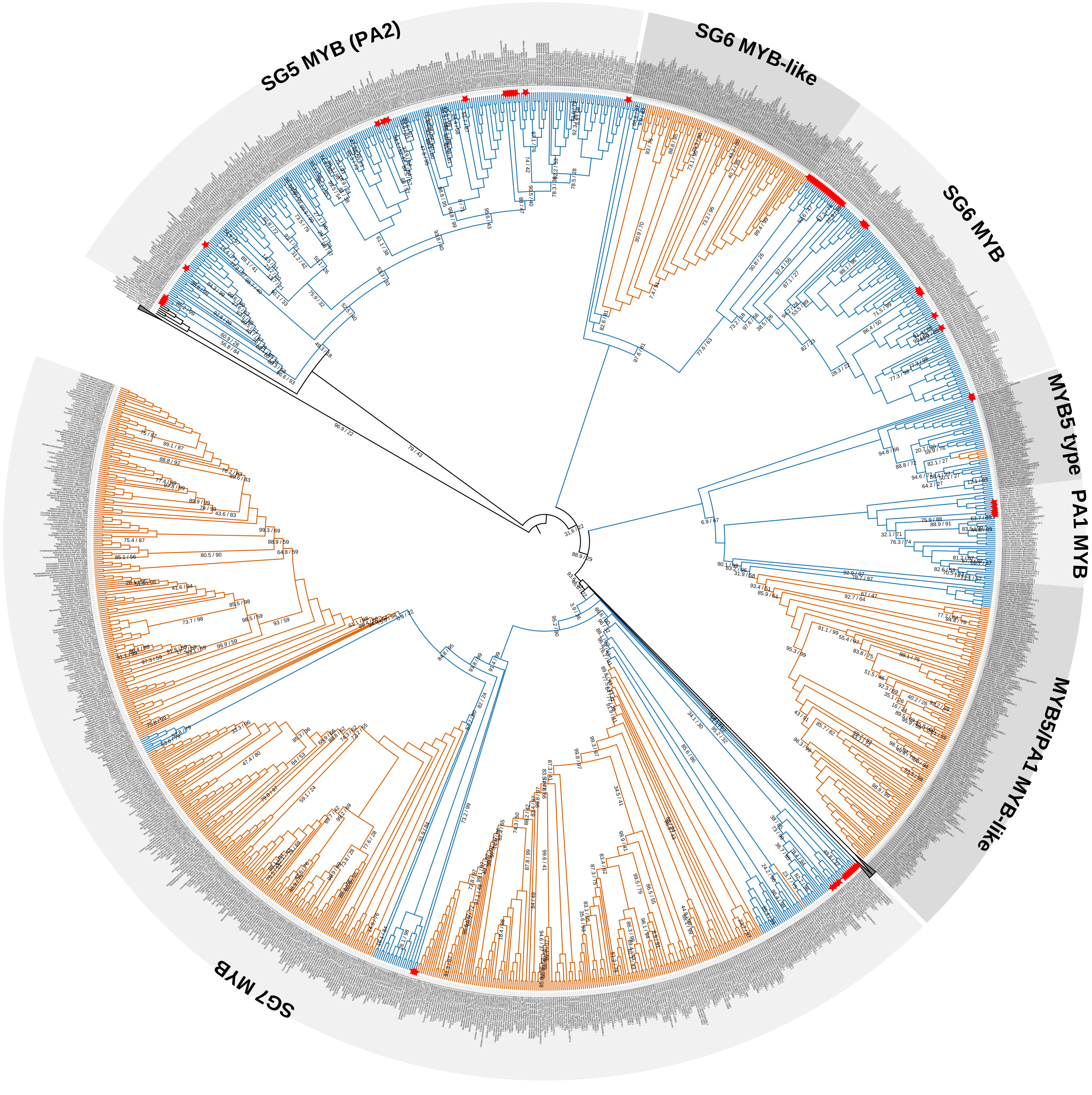


**Fig. S11a: Codon-based maximum-likelihood phylogenetic tree of MYB sequences constructed with MAFFT and IQ-TREE.**

The best-fit substitution model selected by Model-Finder was GTR+F+ASC+R10. Cucurbitaceae sequences are shown in orange branches, and non-Cucurbitaceae sequences shown in blue. Red stars at the tip of terminal branches indicate functional enzymatic sequences identified in previous studies. Numbers above nodes represent Shimodaira-Hasegawa approximate likelihood ratio test (SH-aLRT)support (%) / ultrafast bootstrap support (%) based on 1000 replicates. Only percentages below 100 are shown. MYB, myeloblastosis; SG, subgroup; PA MYB, proanthocyanidin MYB
