## Supplementary material for "Out of the blue: Family-wide loss of anthocyanin biosynthesis in Cucurbitaceae": Fig. S12a

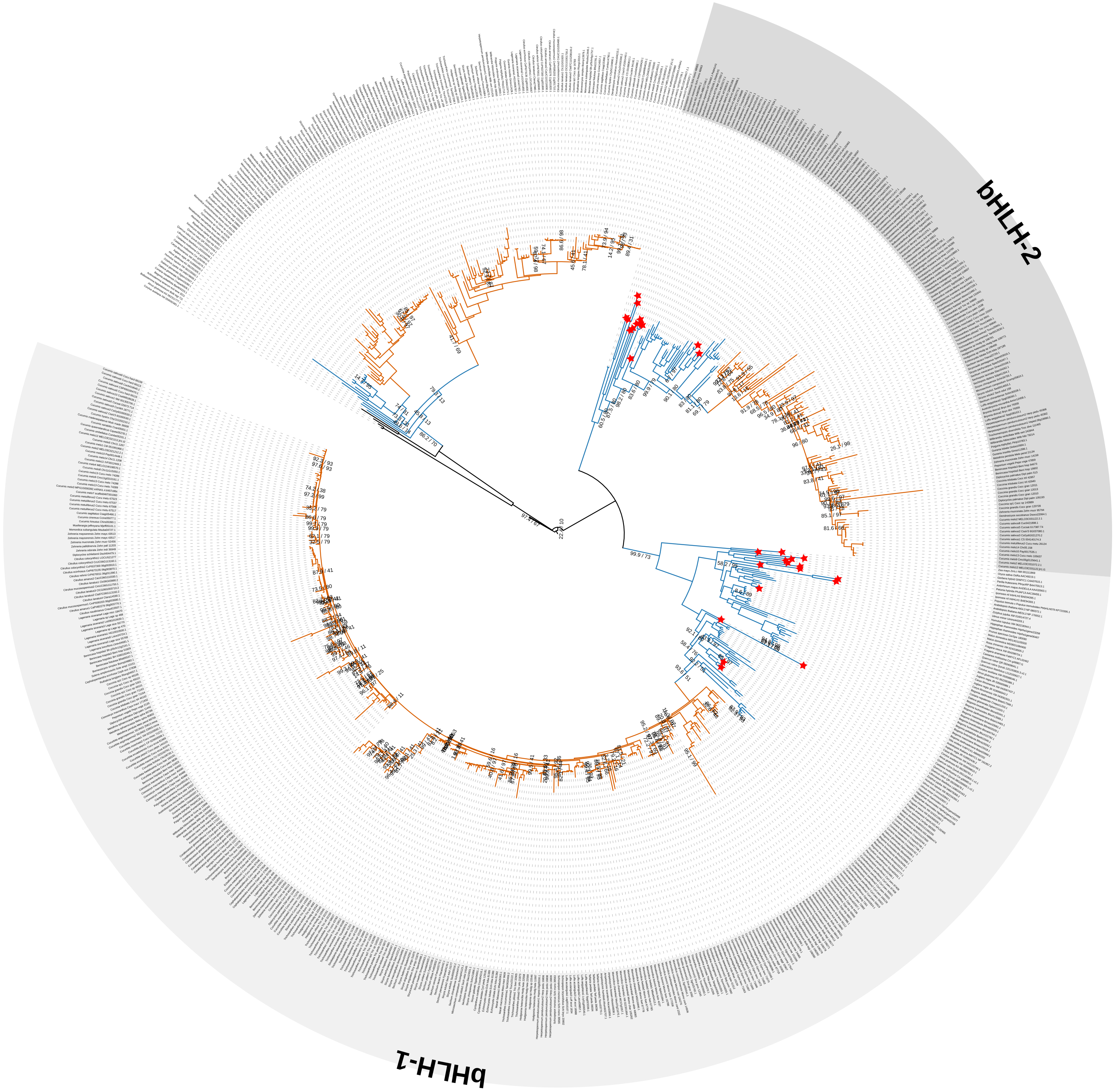


**Fig. S12a: Codon-based maximum-likelihood phylogenetic tree of flavonoid-related bHLH sequences constructed with MAFFT and IQ-TREE.**
Flavonoid-related bHLHs are regulators of anthocyanin and/or proanthocyanidin pathway and are part of the MBW (MYB-bHLH-WD40) complex. bHLH1 is the *Arabidopsis thaliana* MYC1 ortholog clade while bHLH2 is the *A. thaliana* TT8 ortholog clade. The best-fit substitution model selected by Model-Finder was GTR+F+ASC+R10. Cucurbitaceae sequences are shown in orange branches, and non-Cucurbitaceae sequences shown in blue. Red stars at the tip of terminal branches indicate functional enzymatic sequences identified in previous studies. Numbers above nodes represent Shimodaira-Hasegawa approximate likelihood ratio test (SH-aLRT) support (%) / ultrafast bootstrap support (%) based on 1000 replicates. Only percentages below 100 are shown. bHLH, basic helix-loop-helix.
