## Supplementary material for "Out of the blue: Family-wide loss of anthocyanin biosynthesis in Cucurbitaceae": Fig. S13b

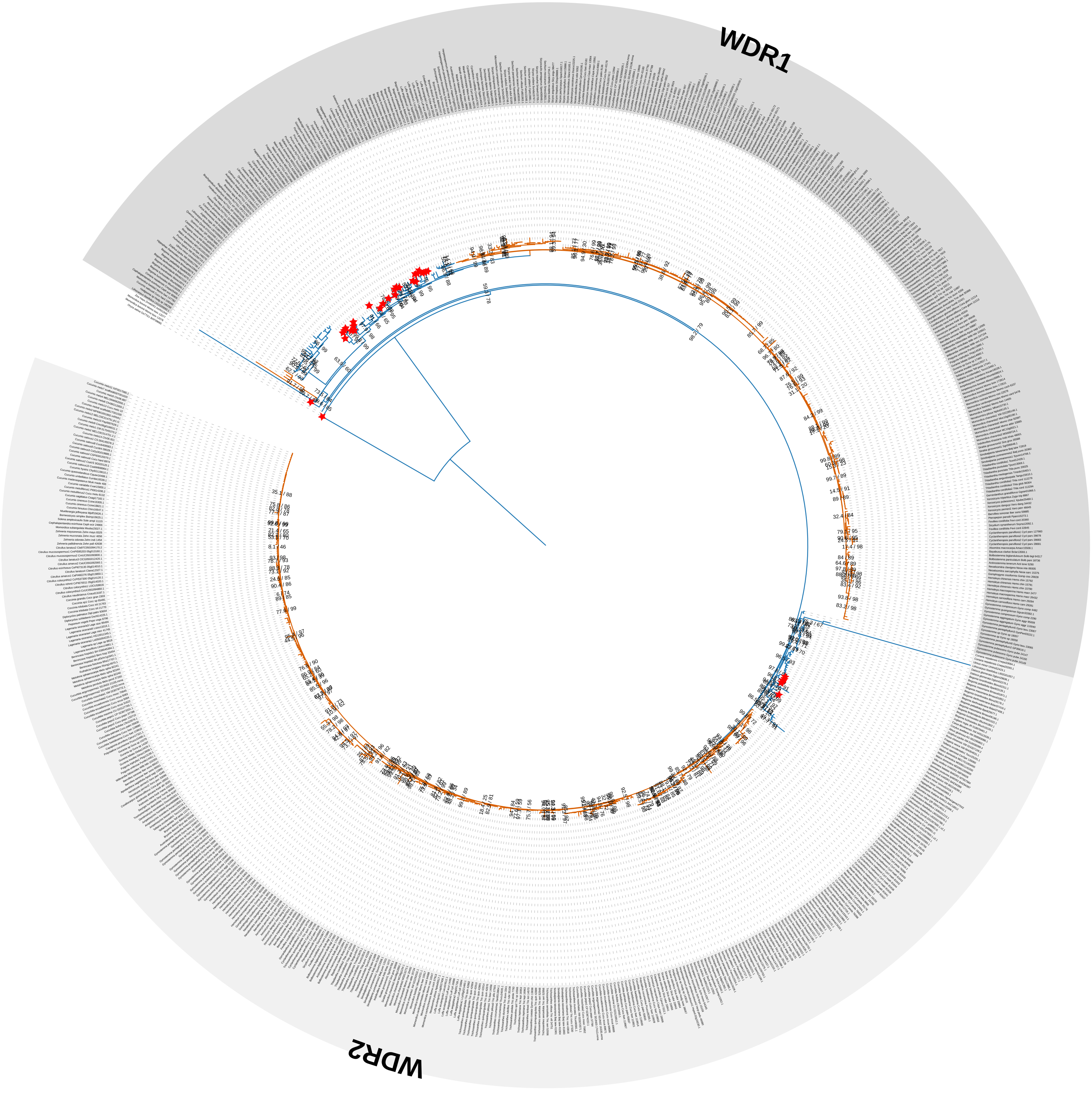


**Fig. S13b: Codon-based maximum-likelihood phylogenetic tree of flavonoid-related WD40 sequences constructed with MUSCLE and IQ-TREE.**

Flavonoid-related WD40s are regulators of anthocyanin and/or proanthocyanidin pathway and are part of the MBW (MYB-bHLH-WD40) complex. WDR1 is the *Arabidopsis thaliana* TTG1 ortholog clade while WDR2 is the *A. thaliana* AN11 ortholog clade. The best-fit substitution model selected by Model-Finder was TIM3+F+R6. Cucurbitaceae sequences are shown in orange branches, and non-Cucurbitaceae sequences shown in blue. Red stars at the start of terminal branch indicate functional enzymatic sequences identified in previous studies. Numbers above nodes represent Shimodaira-Hasegawa approximate likelihood ratio test (SH-aLRT) support (%) / ultrafast bootstrap support (%) based on 1000 replicates. Only percentages below 100 are shown. WDR, WD40-repeat
