## Supplementary material for "Out of the blue: Family-wide loss of anthocyanin biosynthesis in Cucurbitaceae": Fig. S14

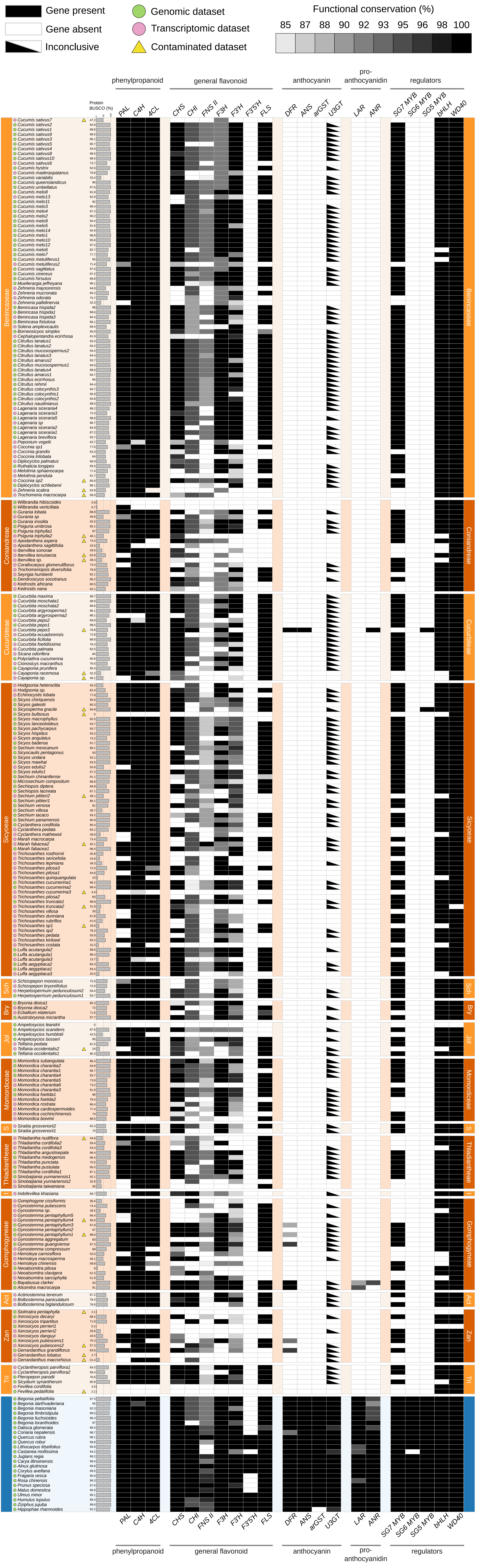


**Fig. S14: Presence-absence matrix of flavonoid biosynthesis genes across all datasets analyzed in this study.** Outgroup species (blue) and Cucurbitaceae species (orange) are annotated by data source, with genomic datasets shown as green circle and transcriptomic datasets as pink circles. Yellow triangles next to dataset names denote datasets identified as contaminated based on Kraken classification. The percentage of conserved BUSCO genes identified in the polypeptide sequences of each dataset is shown as grey horizontal bars. Gene presence, absence, or putative non-functional sequences are shown as black, white, and grey boxes, respectively. Tribe abbreviations: Tri, Triceratieae; Zan, Zanonieae; Act, Actinostemmateae; I, Indofevilleae; S; Siraitieae; Jol, Joliffieae; Bry, Bryonieae; Sch, Schizopeponeae
