## Supplementary material for "Out of the blue: Family-wide loss of anthocyanin biosynthesis in Cucurbitaceae": Fig. S15

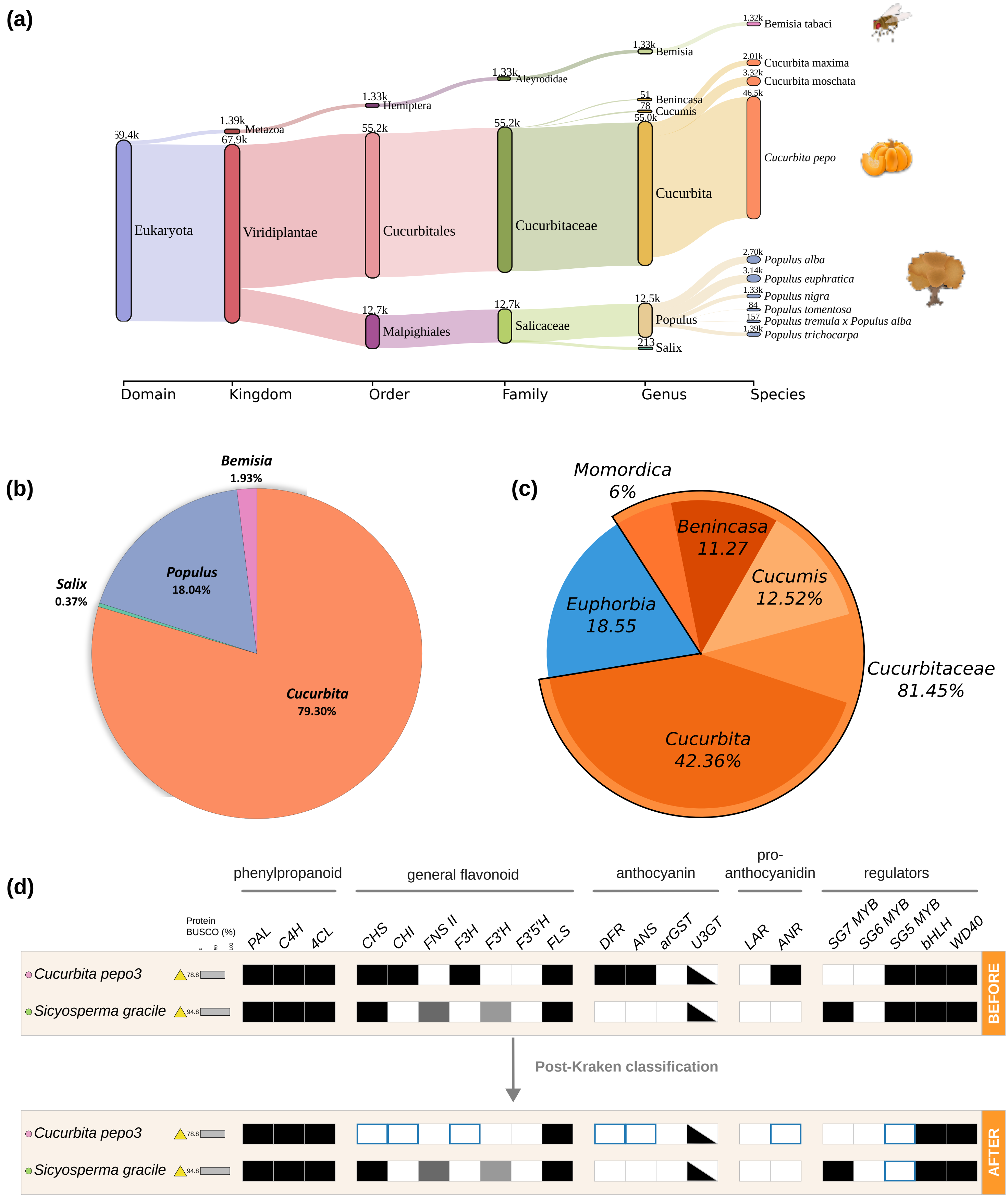


**Fig. S15: Contamination profiles of *Cucurbita pepo*3 and *Sicyosperma gracile***

(a) Sankey diagram showing taxonomic classification of *C. pepo*3 sequences from higher taxonomic ranks to species level.

(b) Pie chart summarizing species-level classification of *C. pepo*3 sequences, highlighting contamination from *Populus* species.

(c) Pie chart summarizing species-level classification of *S. gracile* sequences, highlighting contamination from *Euphorbia* species.

(d) Presence-absence matrix of anthocyanin biosynthesis genes in contaminated plant datasets compared with the matrix after contamination check using Kraken. Gene presence is shown as coloured boxes (black/grey) and absence as white boxes. Sequences assigned to non-Cucurbitaceae taxa are removed and highlighted with a blue border, indicating their origin from contaminant species (*Populus* in *C. pepo*3 and *Euphorbia* in *S. gracile*).

These results highlight the importance of contamination screening prior to gene presence-absence analyses, particularly when contamination originates from anthocyanin-pigmented lineages, which can lead to false-positive gene identifications.
