## Supplementary material for "Out of the blue: Family-wide loss of anthocyanin biosynthesis in Cucurbitaceae": Fig. S16

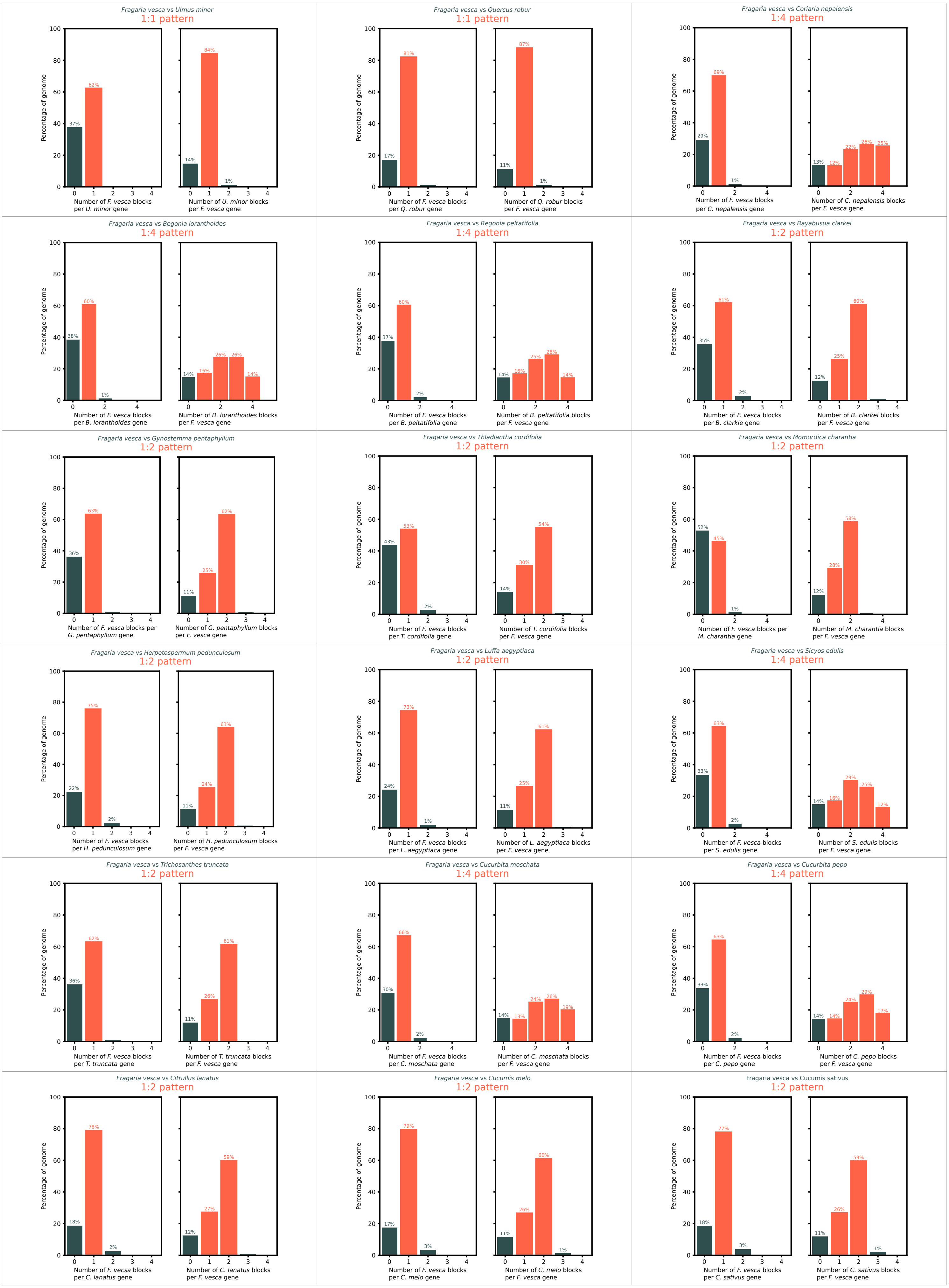


**Fig. S16: Synteny depth plots of reference vs target datasets used in the microsynteny analysis**
*Fragaria vesca* was used as the reference species. All plots were generated using JCVI MCscan.
