## Supplementary material for "Out of the blue: Family-wide loss of anthocyanin biosynthesis in Cucurbitaceae": Fig. S17

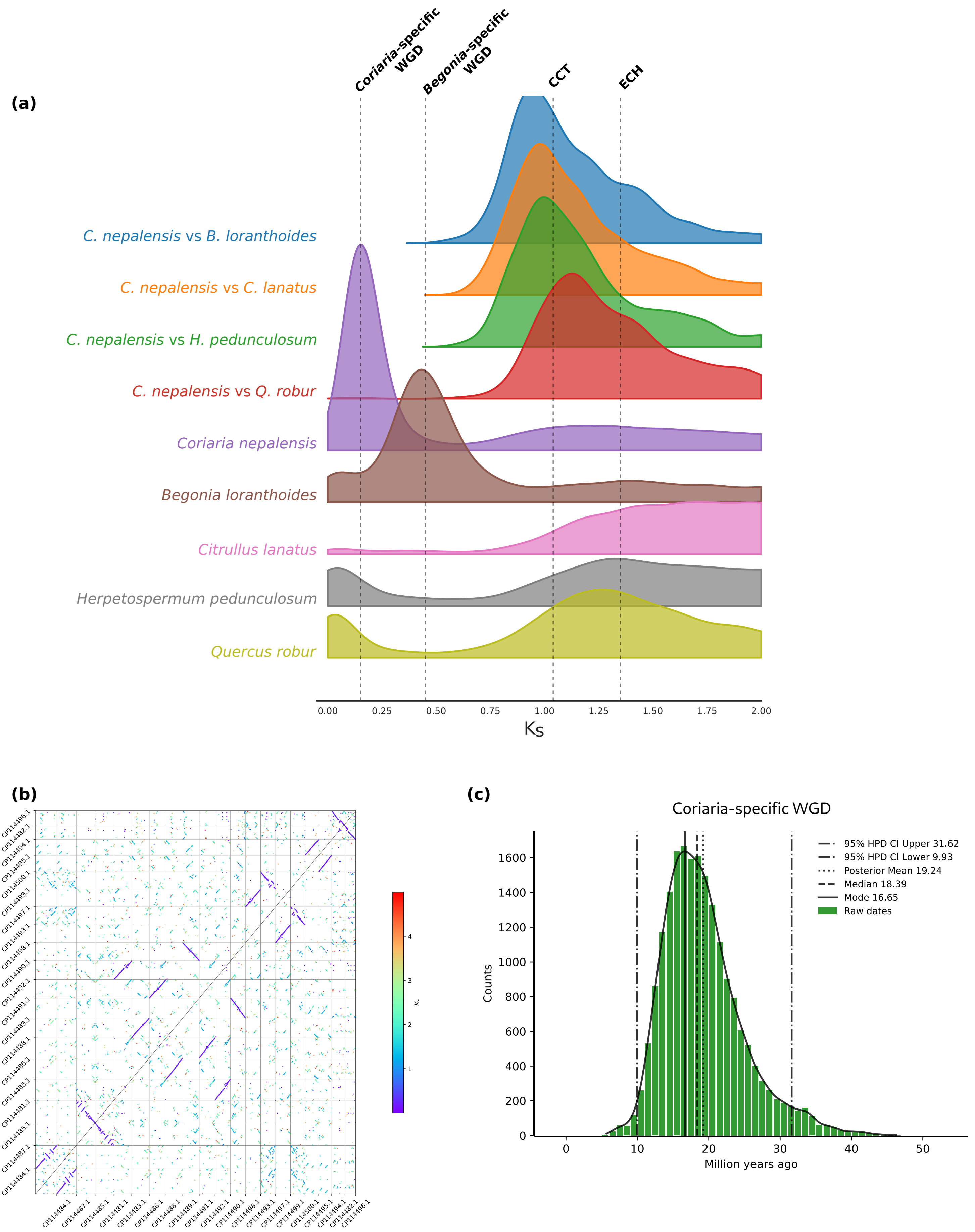


**Fig. S17: Whole Genome Duplication (WGD) detection and dating in *Coriaria nepalensis***
(a) The whole paranome *Ks* (synonymous nucleotide substitution rate) distributions of *C. nepalensis* and closely related species, along with evolutionary rate-corrected divergence events (speciation events) based on orthologs btewеen these species and *C. nepalensis*.
(b) Intraspecific genome homology dotplots for *C. nepalensis*. Anchor pairs are represented as dots coloured by their associated *Ks* values. Axis show genes on corresponding chromosomes. A large number of homologous genes with a low *Ks* value indicates a recent WGD.
(c) Posterior date distribution, overall mean, and 95% Highest Posterior Density (HPD) of the estimated date of the identified *Coriaria*-specific WGD.
ECH: Core eudicot common hexaploidization; CCT: Cucurbitales common tetraploidization
