## Supplementary material for "Out of the blue: Family-wide loss of anthocyanin biosynthesis in Cucurbitaceae": Fig. S18

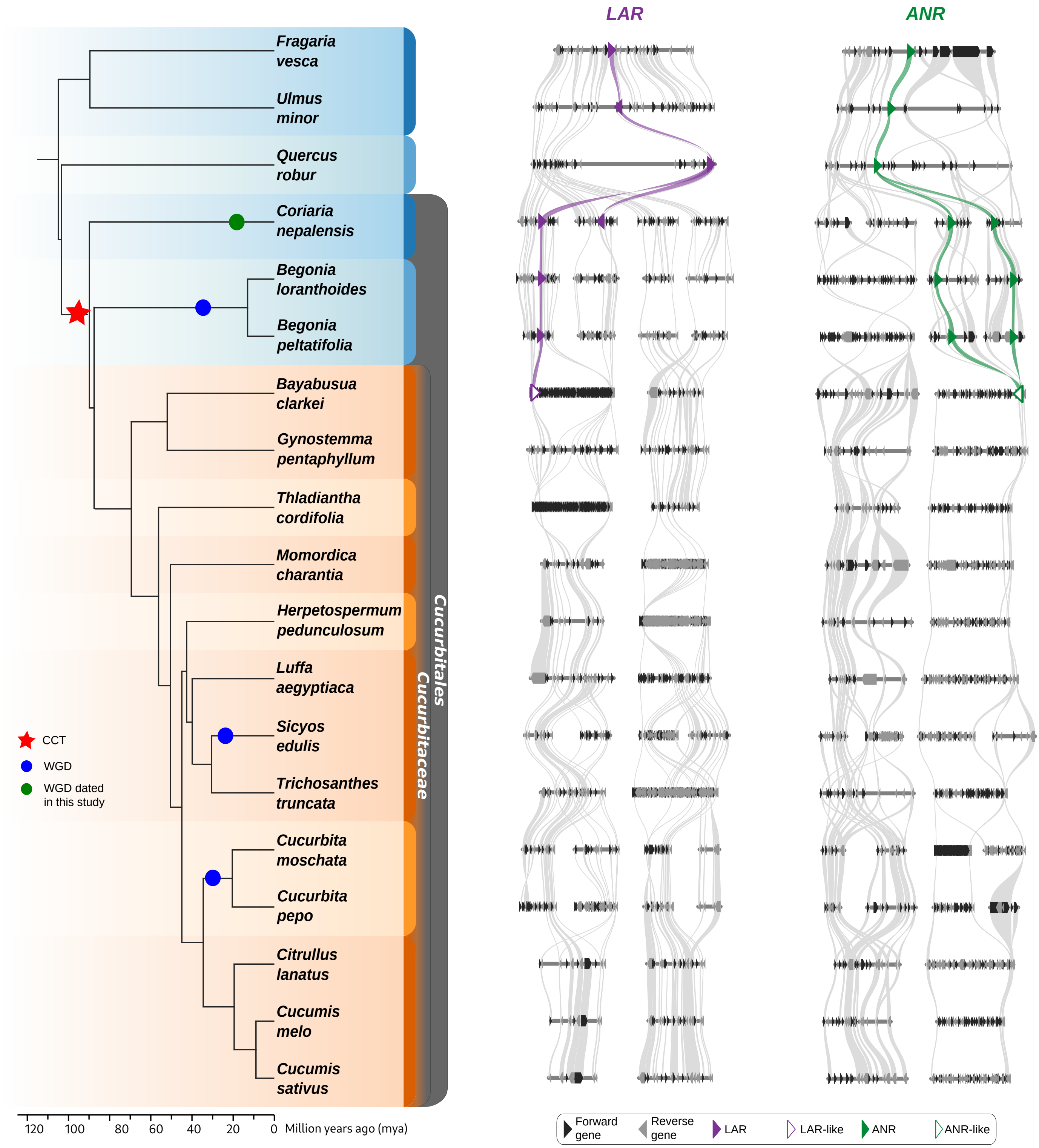


**Fig. S18: Microsynteny around LAR and ANR loci.**
On the left, the species phylogeny is shown. A red star denotes the Cucurbitales common tetraploidization (CCT) event, while blue circles indicate lineage-specific whole-genome duplications (WGDs): the Begoniaceae common tetraploidization (BCT), the Sicyoeae WGD, and the *Cucurbita*-specific polyploidization (CST). A green circle marks the *Coriaria*-specific WGD dated in this study. On the right, conserved syntenic regions for the proanthocyandin biosynthesis genes, *LAR* (purple) and *ANR* (green) are shown. Genes outlines with unfilled coloured borders represent putatively non-functional copies, based on conserved amino acid residue analysis. Other genes are shown in black (forward strand) and grey (reverse strand). The number of syntenic blocks displayed per species reflects the degree of collinearity relative to *Fragaria vesca* and corresponds to the number of polyploidization events in the evolutionary history of each lineage.

*Note: In Begonia species, microsynteny analysis of the ANR locus identified only three syntenic regions corresponding to F. vesca; therefore, only three regions are shown.*
