## Supplementary material for "Out of the blue: Family-wide loss of anthocyanin biosynthesis in Cucurbitaceae": Fig. S19

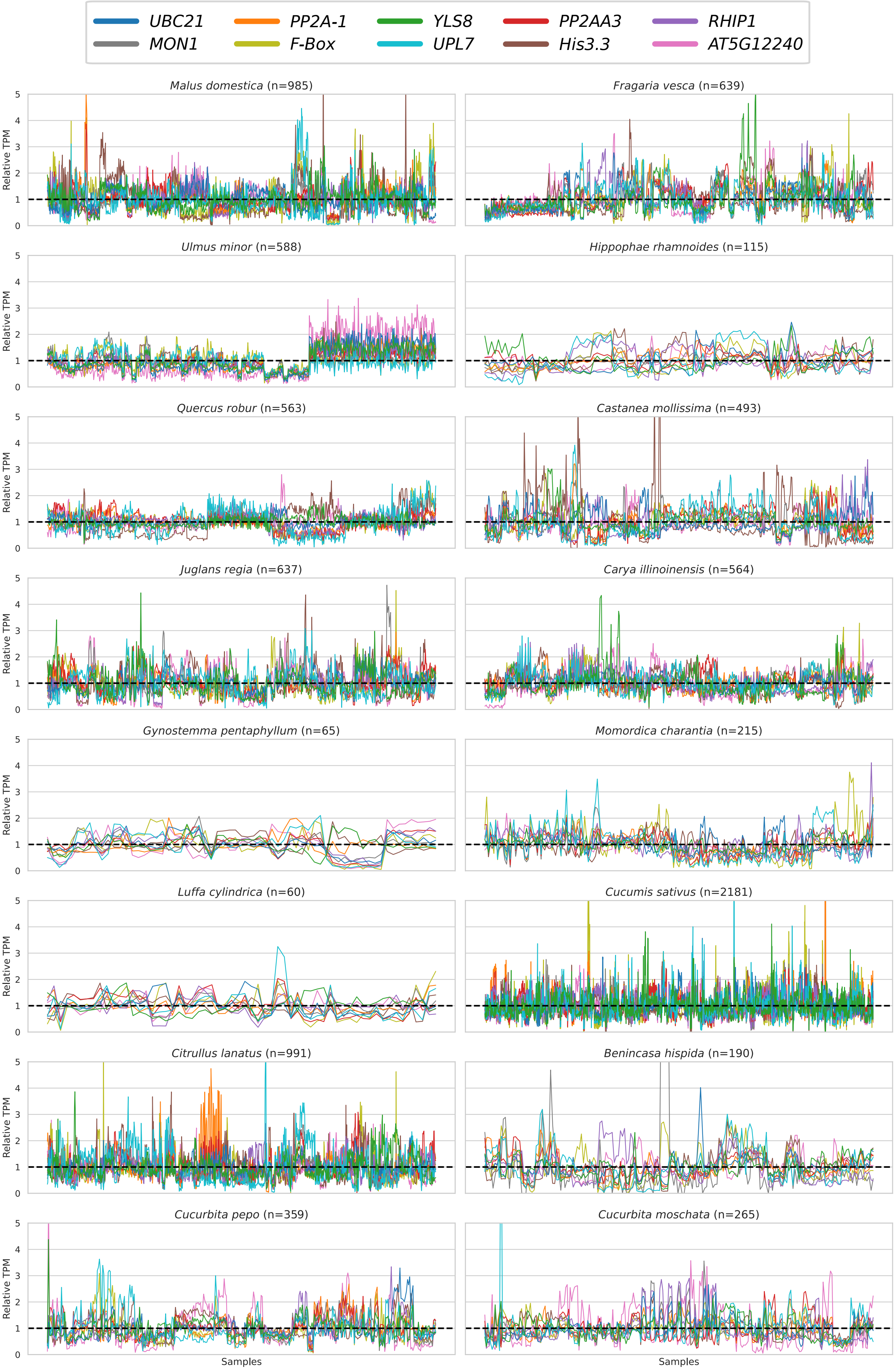


**Fig. S19: Relative expression stability of selected reference genes across 16 plant species.**
For each species, the relative expression of each reference gene, expressed as transcripts per million (TPM) and calculated as the expression in each sample divided by the mean expression across all samples, is shown across all samples. Sample size (n) for each species is indicated above the plots.
