## Supplementary material for "Out of the blue: Family-wide loss of anthocyanin biosynthesis in Cucurbitaceae": Table S1

**Table S1: Candidate reference genes evaluated for expression normalization**
Reference genes previously reported to show stable expression in *Arabidopsis thaliana* were collected from literature and assessed for expression stability across all species and samples analyzed in this study. Genes were classified as *Selected* based on the stability of their transcript abundance (see Methods). Reasons for exclusion are provided where applicable.

| ***A. thaliana* ID** | **Gene Name / Function** | **Gene Symbol** | **Status** | **Reason** | **Reference** |
|---|---|---|---|---|---|
| AT1G13320 | serine/threonine protein phosphatase 2A subunit A3 | *PP2AA3* | Selected | Stable transcript abundance across all samples | Czechowski *et al.*, 2005 |
| AT1G59830 | serine/threonine protein phosphatase PP2A-1 catalytic subunit | *PP2A-1* | Selected | Stable transcript abundance across all samples | Czechowski *et al.*, 2005 |
| AT2G28390 | monensin sensitivity1; SAND family protein | *MON1* | Selected | Stable transcript abundance across all samples | Czechowski *et al.*, 2005 |
| AT3G53090 | ubiquitin-protein ligase 7 | *UPL7* | Selected | Stable transcript abundance across all samples | Czechowski *et al.*, 2005 |
| AT4G26410 | RGS1-HXK1 interacting protein 1 | *RHIP1* | Selected | Stable transcript abundance across all samples | Souček *et al.*, 2017 |
| AT4G40030 | histone variant H3 | His3.3 | Selected | Stable transcript abundance across all samples | Ferreira *et al.*, 2023 |
| AT5G08290 | yellow-leaf-specific gene 8 | *YLS8* | Selected | Stable transcript abundance across all samples | Czechowski *et al.*, 2005 |
| AT5G12240 | octanoyltransferase | *Expressed -protein* | Selected | Stable transcript abundance across all samples | Czechowski *et al.*, 2005 |
| AT5G15710 | galactose oxidase/kelch repeat superfamily protein | *Fbox* | Selected | Stable transcript abundance across all samples | Czechowski *et al.*, 2005 |
| AT5G25760 | ubiquitin-conjugating enzyme 21 | *UBC21* | Selected | Stable transcript abundance across all samples | Czechowski *et al.*, 2005 |
| AT1G58050 | helicase domain-containing protein | *Helicase* | Excluded | Ortholog found; comparatively less stable expression across all samples than the selected ones | Czechowski *et al.*, 2005 |
| AT2G32170 | methyltransferase gene | *EXPRS* | Excluded | Ortholog found; comparatively less stable expression across all samples than the selected ones | Czechowski *et al.*, 2005 |
| AT5G46630 | adaptor protein-2 mu-adaptin | *AP2M* | Excluded | Ortholog found; comparatively less stable expression across all samples than the selected ones | Czechowski *et al.*, 2005 |
| AT4G27960 | ubiquitin-conjugating enzyme 9 | *UBC9* | Excluded | Ortholog found; comparatively less stable expression across all samples than the selected ones | Czechowski *et al.*, 2005 |
| AT4G33380 | adenosine tRNA methylthiotransferase | *Expressed -protein* | Excluded | Ortholog found; comparatively less stable expression across all samples than the selected ones | Czechowski *et al.*, 2005 |
| AT4G34270 | TAP42 interacting protein of 41 kda | *TIP41* | Excluded | Ortholog found; comparatively less stable expression across all samples than the selected ones | Czechowski *et al.*, 2005 |
| AT1G50010 | tubulin alpha-2 chain | *TUA2* | Excluded | Ortholog not identified in *C. illinoinensis* and *J. regia* | Ferreira *et al.*, 2023 |
| AT3G02470 | S-adenosylmethionine decarboxylase | *SAMDC* | Excluded | Ortholog not identified in *L. aegyptiaca* | Ferreira *et al.*, 2023 |
| AT3G18780 | actin 2 | *ACT2* | Excluded | Ortholog not identified in *F. vesca* | Souček *et al.*, 2017 |
| AT4G05320 | ubiquitin 10 | *UBQ10* | Excluded | Ortholog not identified in *F. vesca*, *H. rhamnoides*, and *Q. robur* | Souček *et al.*, 2017 |
| AT1G62930 | RNA processing factor 3 | *RPF3* | Excluded | Ortholog not identified in *L. aegyptiaca*, *M. charantia*, *C. sativus*, *B. hispida*, *C. lanatus*, *C. pepo*, *C. moschata* | Czechowski *et al.*, 2005 |
| AT4G38070 | transcription factor bHLH131-like protein | *bHLH131* | Excluded | Ortholog not identified in *U. minor* and *G. pentaphyllum* | Czechowski *et al.*, 2005 |
| AT5G09810 | actin 7 | *ACT7* | Excluded | Ortholog not identified in *H. rhamnoides* | Ferreira *et al.*, 2023 |
| AT1G30950 | unusual floral organs | *UFO* | Excluded | Ortholog not identified in *L. aegyptiaca* | Hong *et al.*, 2010 |
| AT5G55840 | PPR superfamily protein | *PPR* | Excluded | Ortholog not identified in *U. minor* and *H. rhamnoides* | Czechowski *et al.*, 2005 |
| AT5G60390 | elongation factor 1-alpha 4 | *EF-1α* | Excluded | Failed to identify clear ortholog clade | Souček *et al.*, 2017 |
| AT5G62690 | tubulin beta chain 2 | *TUB2* | Excluded | Failed to identify clear ortholog clade | Hong *et al.*, 2010 |
| AT1G07920 | elongation factor 1-alpha 3 | *EF-1α3* | Excluded | Failed to identify clear ortholog clade | Souček *et al.*, 2017 |
| AT1G13440 | glyceraldehyde-3-phosphare dehydrogenase | *GAPDH* | Excluded | Failed to identify clear ortholog clade | Czechowski *et al.*, 2005 |
| AT1G14320 | suppressor of ACAULIS 52 | *SAC52* | Excluded | Failed to identify clear ortholog clade | Ferreira *et al.*, 2023 |
| AT3G01150 | polypyrimidine tract-binding protein | *PTB* | Excluded | Failed to identify clear ortholog clade | Czechowski *et al.*, 2005 |
| AT4G36800 | RUB1 conjugating enzyme 1 | *RCE1* | Excluded | Failed to identify clear ortholog clade | Ferreira *et al.*, 2023 |
