## Supplementary material for "Out of the blue: Family-wide loss of anthocyanin biosynthesis in Cucurbitaceae": Table S2

**Table S2: Reference gene orthologs used for expression normalisation across species**
For each of the ten selected reference genes, corresponding orthologous gene models used for expression analysis are listed for each species.

| | ***PP2AA3*** | ***PP2A-1*** | ***MON1*** | ***UPL7*** | ***RHIP1*** | ***His3.3*** | ***YLS8*** | ***Expressed-protein*** | ***Fbox*** | ***UBC21*** |
|---|---|---|---|---|---|---|---|---|---|---|
| ***Arabidopsis thaliana*** | **AT1G13320** | **AT1G59830** | **AT2G28390** | **AT3G53090** | **AT4G26410** | **AT4G40030** | **AT5G08290** | **AT5G12240** | **AT5G15710** | **AT5G25760** |
| ***Malus domestica*** | MD13G1044500, MD16G1045400, MD06G1214700, MD14G1225600 | MD13G1083600, MD16G1083000, MD14G1189900, MD06G1183800 | MD03G1118200, MD11G1136600 | MD12G1101300, MD14G1095600 | MD11G1001100 | MD15G1320600, MD15G1373700 | MD16G1003600, MD13G1003900 | MD11G1174500 | MD09G1076600 | MD08G1116800, MD15G1096300 |
| ***Fragaria vesca*** | XM_004300486.1, XM_004297079.1 | XM_004299571.1, XM_004297640.1 | XM_004293004.1 | XM_004303006.1 | XM_004294885.1 | XM_004299624.1, XM_004291779.1, XM_004288352.1, XM_004288116.1, XM_004287165.1 | XM_004293850.1, XM_004297683.1 | XM_004291484.1 | XM_004304138.1 | XM_004291147.1 |
| ***Ulmus minor*** | Umino36473.1, Umino46550.1 | Umino45791.1, Umino35680.1 | Umino28814.1 | Umino50448.1 | Umino18891.1 | Umino39484.1, Umino05842.1 | Umino51084.1 | Umino02358.1 | Umino04777.1 | Umino06817.1 |
| ***Hippophae rhamnoides*** | Hiprha1gene00320 | Hiprha1gene16857, Hiprha1gene18525, Hiprha1gene25511, Hiprha1gene30501 | Hiprha1gene24930, Hiprha1gene29846 | Hiprha1gene27518 | Hiprha1gene16323 | Hiprha1gene30621, Hiprha1gene29044, Hiprha1gene27419, Hiprha1gene04942, Hiprha1gene04589 | Hiprha1gene18545 | Hiprha1gene27925 | Hiprha1gene21641, Hiprha1gene20154 | Hiprha1gene15154, Hiprha1gene14277 |
| ***Quercus robur*** | QR_050385359.1, QR_050402249.1 | QR_050400796.1, QR_050420939.1 | QR_050430911.1, QR_050388804.1 | QR_050387415.1 | QR_050421740.1 | QR_050436377.1, QR_050435275.1, QR_050411659.1 | QR_050383967.1 | QR_050427206.1 | QR_050414474.1 | QR_050409355.1 |
| ***Castanea mollisima*** | Cm_g15912.t1 | Cm_g6662.t1, Cm_g5221.t1 | Cm_g21920.t1 | Cm_g49605.t1 | Cm_g49767.t1 | Cm_g11307.t1 | Cm_g20732.t1 | Cm_g4925.t1 | Cm_g17260.t1 | Cm_g2572.t1 |
| ***Juglans regia*** | JR_035685846.1, JR_018953794.2, JR_018974404.2 | JR_018963562.2, JR_035684549.1 | JR_018963979.2, JR_035690214.1 | JR_018988558.2 | JR_018981994.2 | JR_018993457.2, JR_018992219.2, JR_018978364.2, JR_018965606.2 | JR_018974444.2, JR_018961779.2 | JR_035690453.1 | JR_018961389.2 | JR_035693748.1 |
| ***Carya illinoinensis*** | XM_043109563.1, XM_043105771.1 | XM_043128385.1, XM_043124679.1 | XM_043099773.1, XM_043103013.1 | XM_043086155.1 | XM_043101211.1 | XM_043133397.1, XM_043120035.1, XM_043115527.1, XM_043104386.1 | XM_043122040.1, XM_043094639.1 | XM_043103934.1 | XM_043114513.1 | XM_043097133.1 |
| ***Gynostemma pentaphyllum2*** | GP26543.1, GP39264.1, GP39191.1 | GP17806.1, GP24603.1, GP24730.1 | GP03979.1, GP03839.1 | GP08413.1 | GP22475.1, GP22360.1 | GP18640.1, GP36564.1, GP36565.1 | GP26132.1 | GP35933.1, GP36792.1 | GP02000.1 | GP18502.1 |
| ***Momordica charantia2*** | Moc09g37880.1 | Moc09g39880.1, Moc08g32520.1 | Moc03g22040.1 | Moc05g30760.1 | Moc06g09890.1 | Moc08g02280.1 | Moc08g39760.1, Moc09g34770.1 | Moc02g01960.1 | Moc04g37150.1 | Moc08g03380.1 |
| ***Luffa aegyptiaca1*** | Lcy13g004410.1 | Lcy01g020090.1, Lcy13g002280.1 | Lcy10g004280.1 | Lcy06g020730.1 | Lcy03g010700.1 | Lcy11g009740.1 | Lcy13g008340.1 | Lcy07g017940.1 | Lcy09g003070.1 | Lcy01g011310.1 |
| ***Cucumis sativus1*** | CS_011657465.2 | CS_004135164.3, CS_004146712.3 | CS_004139400.3 | CS_031882877.1 | CS_004144016.3 | CS_031889847.1, CS_031889801.1, CS_031889800.1, CS_004152577.3, CS_004146827.3 | CS_031881493.1, CS_004143570.3 | CS_004140136.3 | CS_031885920.1 | CS_004150235.3 |
| ***Citrullus lanatus2*** | Cla97C05G104770.1 | Cla97C05G106960.1, Cla97C08G151900.1 | Cla97C01G012740.1 | Cla97C10G202640.2 | Cla97C03G064890.2 | Cla97C08G154270.1 | Cla97C02G038590.1, Cla97C02G039800.1 | Cla97C02G048780.1 | Cla97C11G211810.2 | Cla97C08G155510.2 |
| ***Benincasa hispida1*** | BH_039036531.1 | BH_039035994.1, BH_039028886.1 | BH_039038527.1 | BH_039048935.1 | BH_039024044.1 | BH_039030074.1 | BH_039046947.1 | BH_039044486.1 | BH_039034704.1 | BH_039030857.1 |
| ***Cucurbita pepo1*** | CP_023680913.1, CP_023686598.1 | CP_023658059.1, CP_023690273.1, CP_023681854.1 | CP_023664294.1 | CP_023683460.1 | CP_023661056.1 | CP_023675962.1, CP_023691263.1, CP_023697253.1 | CP_023657340.1, CP_023676953.1, CP_023687468.1 | CP_023676616.1 | CP_023657641.1, CP_023693801.1 | CP_023680983.1, CP_023689814.1 |
| ***Cucurbita moschata1*** | XM_023101179.1, XM_023089599.1 | XM_023095857.1, XM_023090726.1, XM_023101151.1 | XM_023109205.1 | XM_023091022.1 | XM_023105465.1 | XM_023073716.1, XM_023086482.1, XM_023106719.1 | XM_023077391.1, XM_023101318.1, XM_023086844.1, XM_023072211.1 | XM_023102091.1 | XM_023084181.1, XM_023109268.1 | XM_023095212.1, XM_023106552.1 |
