## Supplementary material for "Out of the blue: Family-wide loss of anthocyanin biosynthesis in Cucurbitaceae": Table S3

**Table S3: Genes used for expression analyses of carotenoid biosynthesis genes in outgroups and Cucurbitaceae**

For each species, gene IDs corresponding to carotenoid biosynthesis enzymes are listed. Functional conservation (FC, %) indicates the percentage of conserved amino acid residues required for the annotated enzyme function.

*For CrtISO, expression values of *CrtISO1* and *CrtISO2* gene candidates were added to derive total CrtISO expression shown in Fig. 5.

| | *Z-ISO* | | *ZDS* | | *CrtISO1** | | *CrtISO2** | | *LCYE* | | *LCYB* | | *CYP97A* | | *BCH* | | *VDE* | | *ZEP* | | *NXS* | |
|---|---|---|---|---|---|---|---|---|---|---|---|---|---|---|---|---|---|---|---|---|---|---|
| | Gene ID | FC (%) | Gene ID | FC (%) | Gene ID | FC (%) | Gene ID | FC (%) | Gene ID | FC (%) | Gene ID | FC (%) | Gene ID | FC (%) | Gene ID | FC (%) | Gene ID | FC (%) | Gene ID | FC (%) | Gene ID | FC (%) |
| ***Malus domestica*** | MD04G1028100 | 100 | MD04G1220900 | 100 | MD14G1044400 | 100 | MD08G1243700 | 100 | MD02G1083500 | 100 | MD00G1049000 | 100 | MD04G1239800 | 100 | MD01G1208300 | 100 | MD12G1255200 | 100 | MD02G1172400 | 100 | MD06G1045700 | 100 |
| | | | MD12G1237300 | 100 | | | | | | | | | MD12G1257000 | 100 | MD07G1278900 | 100 | | | MD15G1284500 | 100 | MD00G1132400 | 100 |
| | | | | | | | | | | | | | | | MD06G1089200 | 100 | | | | | MD09G1253000 | 100 |
| ***Fragaria vesca*** | XM_004297980.1 | 100 | XM_004301977.1 | 100 | XM_004303334.1 | 100 | XM_004301509.1 | 100 | XM_004287534.1 | 100 | XM_004303559.1 | 100 | XM_004302087.1 | 100 | XM_004308006.1 | 100 | XM_004302077.1 | 100 | XM_004306336.1 | 93.3 | XM_004300264.1 | 100 |
| | | | | | | | | | | | | | | | XM_004300765.1 | 100 | | | | | | |
| ***Ulmus minor*** | Umino51417.1 | 100 | Umino29712.1 | 100 | Umino49671.1 | 100 | Umino30776.1 | 100 | Umino33628.1 | 100 | Umino49479.1 | 100 | Umino30099.1 | 100 | Umino13563.1 | 100 | Umino30042.1 | 100 | Umino38793.1 | 100 | Umino04414.1 | 100 |
| | | | Umino29693.1 | 100 | | | | | | | | | | | Umino44146.1 | 100 | | | | | | |
| ***Hippophae rhamnoides*** | Hiprha1gene09348 | 100 | Hiprha1gene16000 | 100 | Hiprha1gene17015 | 100 | Hiprha1gene16344 | 100 | Hiprha1gene10883 | 100 | Hiprha1gene27597 | 100 | Hiprha1gene26199 | 100 | Hiprha1gene05693 | 100 | Hiprha1gene26091 | 100 | Hiprha1gene15592 | 100 | Hiprha1gene22818 | 100 |
| | | | | | | | | | Hiprha1gene03401 | 100 | | | | | | | | | | | Hiprha1gene00108 | 100 |
| ***Quercus robur*** | QR_050399978.1 | 100 | QR_050419023.1 | 100 | QR_050388536.1 | 100 | QR_050416925.1 | 100 | QR_050414224.1 | 100 | QR_050389511.1 | 100 | QR_050386579.1 | 100 | QR_050397113.1 | 100 | QR_050386520.1 | 100 | QR_050435776.1 | 100 | QR_050413833.1 | 100 |
| | | | | | | | QR_050416927.1 | 100 | QR_050414223.1 | 100 | | | QR_050386575.1 | 100 | QR_050410144.1 | 100 | QR_050386525.1 | 100 | | | | |
| | | | | | | | | | | | | | QR_050386572.1 | 100 | | | QR_050386529.1 | 100 | | | | |
| ***Castanea mollissima*** | Cm_g24958.t1 | 100 | Cm_g17152.t1 | 100 | Cm_g36140.t1 | 100 | Cm_g28965.t1 | 100 | Cm_g21153.t1 | 100 | Cm_g45431.t1 | 100 | Cm_g3450.t1 | 100 | Cm_g34297.t1 | 100 | Cm_g2015.t1 | 100 | Cm_g19348.t1 | 100 | Cm_g40138.t2 | 100 |
| | | | | | | | | | | | Cm_g36967.t1 | 100 | | | Cm_g35921.t1 | 100 | | | | | Cm_g40138.t1 | 100 |
| ***Juglans regia*** | JR_018972132.2 | 100 | JR_018990301.2 | 100 | JR_018993267.2 | 100 | JR_018955462.2 | 100 | JR_018990653.2 | 100 | JR_018961643.2 | 100 | JR_018962194.2 | 100 | JR_018954124.2 | 100 | JR_018971416.2 | 100 | JR_018989429.2 | 100 | JR_018959943.2 | 100 |
| | | | | | | | JR_018955463.2 | 100 | | | | | | | JR_018983068.2 | 100 | | | | | | |
| | | | | | | | | | | | | | | | JR_018997000.2 | 100 | | | | | | |
| ***Carya illinoinensis*** | XM_043115075.1 | 100 | XM_043133515.1 | 100 | XM_043084812.1 | 100 | XM_043127913.1 | 100 | XM_043121056.1 | 100 | XM_043086101.1 | 100 | XM_043132643.1 | 100 | XM_043126582.1 | 100 | XM_043131704.1 | 100 | XM_043118141.1 | 100 | XM_043121614.1 | 100 |
| | | | | | | | | | | | | | | | XM_043129314.1 | 100 | XM_043131705.1 | 100 | | | | |
| | | | | | | | | | | | | | | | XM_043091779.1 | 100 | | | | | | |
| ***Gynostemma pentaphyllum2*** | GP18982.1 | 100 | GP23721.1 | 100 | GP08653.1 | 100 | GP36474.1 | 100 | GP11616.1 | 100 | GP14085.1 | 100 | GP36538.1 | 100 | GP35469.1 | 100 | GP11374.1 | 100 | GP39143.1 | 100 | GP15472.1 | 100 |
| | | | GP23785.1 | 100 | | | | | | | | | | | GP20560.1 | 100 | | | GP39222.1 | 100 | GP14998.1 | 100 |
| ***Momordica charantia2*** | Moc05g08940.1 | 100 | Moc06g26840.1 | 100 | Moc05g29080.1 | 100 | Moc10g05510.1 | 100 | Moc08g45730.1 | 100 | Moc01g17190.1 | 100 | Moc10g06010.1 | 100 | Moc10g01250.1 | 100 | Moc08g38440.1 | 100 | Moc02g13110.1 | 100 | Moc08g12330.1 | 100 |
| | | | | | | | | | | | | | | | Moc06g38230.1 | 100 | | | | | | |
| ***Luffa aegyptiaca1*** | Lcy10g009130.1 | 100 | Lcy03g004530.1 | 100 | Lcy06g018940.1 | 91.6 | Lcy11g011320.1 | 100 | Lcy08g018900.1 | 100 | Lcy04g005840.1 | 100 | Lcy11g010510.1 | 100 | Lcy11g001300.1 | 100 | Lcy08g010620.1 | 100 | Lcy07g004710.1 | 100 | Lcy02g001400.1 | 100 |
| | | | | | | | | | | | | | | | Lcy02g013470.1 | 100 | | | | | | |
| ***Cucumis sativus1*** | CS_004152053.3 | 100 | CS_004142474.3 | 100 | CS_004136487.3 | 100 | CS_011652269.2 | 100 | CS_004141124.3 | 100 | CS_004150713.3 | 100 | CS_004133705.3 | 100 | CS_004140710.3 | 100 | NM_001305728.1 | 100 | NM_001305784.1 | 100 | CS_004145812.3 | 100 |
| | | | | | | | | | | | | | | | CS_004143925.3 | 100 | | | | | CS_031882497.1 | 100 |
| | | | | | | | | | | | | | | | | | | | | | CS_031882498.1 | 100 |
| ***Citrullus lanatus2*** | Cla97C07G142740.2 | 100 | Cla97C06G118930.2 | 100 | Cla97C10G200950.2 | 100 | Cla97C10G190930.2 | 100 | Cla97C11G208040.2 | 100 | Cla97C04G070940.1 | 100 | Cla97C10G191610.1 | 100 | Cla97C05G090480.1 | 100 | Cla97C11G216330.1 | 100 | Cla97C02G038200.1 | 100 | Cla97C05G093720.2 | 100 |
| | | | | | | | | | | | | | | | Cla97C01G002480.2 | 100 | | | | | | |
| ***Benincasa hispida1*** | BH_039026618.1 | 100 | BH_039023482.1 | 100 | BH_039047825.1 | 100 | BH_039049488.1 | 100 | BH_039035238.1 | 100 | BH_039026592.1 | 100 | BH_039050399.1 | 100 | BH_039021520.1 | 100 | BH_039034565.1 | 100 | BH_039045392.1 | 100 | BH_039031262.1 | 100 |
| | | | | | | | | | | | | | | | BH_039051224.1 | 100 | | | | | | |
| ***Cucurbita pepo1*** | CP_023684933.1 | 100 | CP_023674191.1 | 100 | CP_023683621.1 | 100 | CP_023697681.1 | 100 | CP_023682807.1 | 100 | CP_023689700.1 | 100 | CP_023697021.1 | 100 | CP_023696049.1 | 100 | CP_023681009.1 | 100 | CP_023677552.1 | 100 | CP_023671875.1 | 100 |
| | | | | | | | | | CP_023691131.1 | 100 | CP_023661496.1 | 100 | | | CP_023685715.1 | 100 | | | | | CP_023671874.1 | 100 |
| | | | | | | | | | | | | | | | CP_023695199.1 | 100 | | | | | | |
| | | | | | | | | | | | | | | | CP_023695198.1 | 100 | | | | | | |
| ***Cucurbita moschata1*** | XM_023076947.1 | 100 | XM_023077721.1 | 100 | XM_023083737.1 | 100 | XM_023073596.1 | 100 | XM_023081963.1 | 100 | XM_023077230.1 | 100 | XM_023073576.1 | 100 | XM_023089097.1 | 100 | XM_023081828.1 | 100 | XM_023091781.1 | 100 | XM_023079271.1 | 100 |
| | | | | | | | | | XM_023091092.1 | 100 | XM_023069390.1 | 100 | | | XM_023066853.1 | 100 | | | XM_023091780.1 | 100 | XM_023079268.1 | 100 |
| | | | | | | | | | | | | | | | XM_023073255.1 | 100 | | | | | XM_023079270.1 | 100 |
