## Supplementary material for "Out of the blue: Family-wide loss of anthocyanin biosynthesis in Cucurbitaceae": Table S4

### Table S4: Genes used for expression analyses of flavonoid biosynthesis genes in outgroups and Cucurbitaceae

For each species, gene IDs corresponding to flavonoid biosynthesis enzymes are listed. Functional conservation (FC, %) indicates the percentage of conserved amino acid residues required for the annotated enzyme function. '---' indicates gene sequence not detected.

| | ***CHS*** | | ***CHI*** | | ***F3H*** | | ***F3'H*** | | ***FLS*** | | ***DFR*** | | ***ANS*** | | ***arGST*** | | ***LAR*** | | ***ANR*** | |
|---|---|---|---|---|---|---|---|---|---|---|---|---|---|---|---|---|---|---|---|---|
| | Gene ID | FC (%) | Gene ID | FC (%) | Gene ID | FC (%) | Gene ID | FC (%) | Gene ID | FC (%) | Gene ID | FC (%) | Gene ID | FC (%) | Gene ID | FC (%) | Gene ID | FC (%) | Gene ID | FC (%) |
| ***Malus domestica*** | MD04G1003400 | 100 | MD01G1118000 | 100 | MD15G1246200 | 100 | MD06G1201700 | 100 | MD08G1168600 | 100 | MD15G1024100 | 100 | MD03G1001100 | 100 | MD17G1272100 | 100 | MD16G1048500 | 100 | MD05G1335600 | 100 |
| | MD04G1003300 | 100 | MD01G1117800 | 100 | MD02G1132200 | 100 | MD14G1210700 | 100 | MD08G1121600 | 100 | MD08G1028600 | 100 | MD06G1071600 | 100 | | | MD13G1046900 | 100 | | 100 |
| | MD04G1003000 | 100 | MD01G1118100 | | | | | 100 | MD15G1354100 | 100 | | | | | | | MD06G1211400 | 100 | | |
| | MD13G1285100 | 100 | | | | | | | | | | | | | | 100 | | | | |
| ***Fragaria vesca*** | XM_004306495.1 | 100 | XM_004307403.1 | 100 | XM_004287766.1 | 100 | XM_004299260.1 | 100 | XM_004290789.1 | 100 | XM_004291810.1 | 100 | XM_004298672.1 | 100 | XM_004288530.1 | 100 | XM_004297096.1 | 100 | XM_004309614.1 | 100 |
| | XM_004306494.1 | 100 | | | | | | | | | | | | | | | | | | |
| ***Ulmus minor*** | Umino03942.1 | 100 | Umino11916.1 | 100 | Umino37973.1 | 100 | Umino46177.1 | 100 | Umino30799.1 | 100 | Umino05936.1 | 100 | Umino44418.1 | 100 | Umino31194.1 | 100 | Umino36433.1 | 100 | Umino00737.2 | 100 |
| | Umino52268.1 | 100 | | | Umino12489.1 | 100 | | | | | | | | | | | | | | |
| ***Hippophae rhamnoides*** | Hiprha1gene04371 | 100 | Hiprha1gene02313 | 100 | Hiprha1gene24025 | 100 | Hiprha1gene19656 | 100 | Hiprha1gene19276 | 100 | Hiprha1gene25674 | 100 | Hiprha1gene06498 | 100 | --- | | Hiprha1gene24235 | 100 | Hiprha1gene02920 | 100 |
| | Hiprha1gene30592 | 100 | | | Hiprha1gene29544 | 100 | | | Hiprha1gene04820 | 100 | Hiprha1gene11285 | 100 | | | | | Hiprha1gene07946 | 100 | | |
| | Hiprha1gene15431 | 100 | | | | | | | | | | | | | | | Hiprha1gene05323 | 100 | | |
| | Hiprha1gene17498 | 100 | | | | | | | | | | | | | | | | | | |
| ***Quercus robur*** | QR_050414649.1 | 100 | QR_050394358.1 | 100 | QR_050436431.1 | 100 | QR_050400788.1 | 100 | QR_050401176.1 | 100 | QR_050411544.1 | 100 | QR_050408671.1 | 100 | QR_050414417.1 | 100 | QR_050385978.1 | 100 | QR_050432874.1 | 100 |
| | QR_050383154.1 | 100 | | | | | QR_050400789.1 | 100 | | | QR_050411541.1 | 100 | | | | | | | | |
| | | | | | | | QR_050400791.1 | 100 | | | QR_050411542.1 | 100 | | | | | | | | |
| | | | | | | | QR_050400792.1 | 100 | | | | | | | | | | | | |
| | | | | | | | QR_050400790.1 | 100 | | | | | | | | | | | | |
| ***Castanea mollissima*** | Cm_g41904.t1 | 100 | Cm_g26679.t1 | 100 | Cm_g42151.t1 | 100 | --- | 100 | Cm_g26539.t1 | 100 | Cm_g10343.t1 | 100 | Cm_g18792.t1 | 100 | Cm_g44678.t1 | 100 | Cm_g41413.t1 | 95 | Cm_g40265.t1 | 100 |
| | Cm_g6534.t1 | | | | | | | | | | | 100 | | | | 100 | | | | |
| ***Juglans regia*** | JR_018966498.2 | 100 | JR_018972487.2 | 100 | JR_018974079.2 | 100 | JR_018954394.2 | 100 | JR_018965395.2 | 100 | JR_018965609.2 | 100 | JR_018991659.2 | 100 | JR_018954616.2 | 100 | JR_035690843.1 | 100 | JR_018950001.2 | 100 |
| | JR_018971548.2 | 100 | | | | | | | JR_035695146.1 | 100 | | | | | | | JR_035691108.1 | 100 | | |
| | JR_018961711.2 | 100 | | | | | | | JR_018953238.2 | 100 | | | | | | | JR_018980541.2 | 100 | | |
| | | | | | | | | | JR_018953269.2 | 100 | | | | | | | | | | |
| | | | | | | | | | JR_018961611.2 | 100 | | | | | | | | | | |
| ***Carya illinoinensis*** | XM_043130600.1 | 100 | XM_043097661.1 | 100 | XM_043132107.1 | 100 | XM_043093388.1 | 100 | XM_043122765.1 | 100 | XM_043133112.1 | 100 | XM_043093893.1 | 100 | XM_043120910.1 | 100 | XM_043104692.1 | 100 | XM_043087336.1 | 100 |
| | XM_043132918.1 | 100 | | | | | | | XM_043122900.1 | 100 | XM_043133113.1 | 100 | | | | | XM_043109036.1 | 100 | | |
| | XM_043110823.1 | 100 | | | | | | | XM_043118318.1 | 100 | | | | | | | | | | |
| | | | | | | | | | XM_043119140.1 | 100 | | | | | | | | | | |
| | | | | | | | | | XM_043122444.1 | 100 | | | | | | | | | | |
| ***Gynostemma pentaphyllum2*** | GP15028.1 | 100 | GP02930.1 | 100 | GP36709.1 | 100 | GP21533.1 | 97 | GP02556.1 | 100 | GP16318.2 | 89 | --- | | --- | | --- | | --- | |
| | GP06757.1 | 100 | | | | | | | GP29425.1 | 100 | GP16318.1 | 89 | | | | | | | | |
| | GP06947.1 | 100 | | | | | | | | | | | | | | | | | | |
| ***Momordica charantia2*** | Moc08g12970.1 | 100 | Moc04g34280.1 | 95 | Moc10g29680.1 | 100 | Moc06g02700.1 | 83 | Moc04g31040.1 | 100 | --- | | --- | | --- | | --- | | --- | |
| ***Luffa aegyptiaca1*** | Lcy09g021280.1 | 100 | Lcy09g005850.1 | 95 | Lcy11g020370.1 | 100 | Lcy03g015720.1 | 100 | Lcy09g011830.1 | 100 | --- | | --- | | --- | | --- | | --- | |
| ***Cucumis sativus1*** | CS_004145659.3 | 100 | CS_004149769.3 | 95 | CS_004146092.3 | 92 | CS_004138144.3 | 100 | CS_004140814.3 | 100 | --- | | --- | | --- | | --- | | --- | |
| | | | | | CS_004140448.3 | 92 | | | | | | | | | | | | | | |
| ***Citrullus lanatus2*** | Cla97C05G094290.1 | 100 | Cla97C09G181670.1 | 95 | Cla97C10G185310.1 | 100 | Cla97C06G125240.1 | 100 | Cla97C08G148220.1 | 100 | --- | | --- | | --- | | --- | | --- | |
| ***Benincasa hispida1*** | BH_039036415.1 | 100 | BH_039034903.1 | 90 | BH_039020680.1 | 100 | BH_039022770.1 | 100 | BH_039029707.1 | 100 | --- | | --- | | --- | | --- | | --- | |
| | | | | | BH_039040061.1 | 100 | | | | | | | | | | | | | | |
| ***Cucurbita pepo1*** | CP_023672485.1 | 100 | CP_023693934.1 | 95 | CP_023696035.1 | 92 | CP_023673727.1 | 100 | CP_023695380.1 | 100 | --- | | --- | | --- | | --- | | --- | |
| | | | CP_023693933.1 | | CP_023684146.1 | 92 | CP_023673726.1 | 100 | CP_023660628.1 | 100 | | | | | | | | | | |
| ***Cucurbita moschata1*** | XM_023078567.1 | 100 | XM_023082756.1 | 95 | XM_023067406.1 | 100 | XM_023077843.1 | 100 | XM_023108724.1 | 100 | --- | | --- | | --- | | --- | | --- | |
| | | | | | | | | | XM_023108732.1 | 100 | | | | | | | | | | |
| | | | | | | | | | XM_023082614.1 | 100 | | | | | | | | | | |
